# Integration of APSIM and PROSAIL models to develop more precise radiometric estimation of crop traits using deep learning

**DOI:** 10.1101/2021.02.02.429471

**Authors:** Qiaomin Chen, Bangyou Zheng, Tong Chen, Scott Chapman

**Affiliations:** School of Agriculture and Food Sciences, The University of Queensland, St Lucia, 4067, QLD, Australia; CSIRO Agriculture and Food, Queensland Biosciences Precinct 306 Carmody Road, St Lucia, 4067, QLD, Australia; School of Information Technology and Electrical Engineering, The University of Queensland, St Lucia, 4067, QLD, Australia

**Keywords:** model integration, neural network, hyperspectral data, variable retrieval

## Abstract

A major challenge for the estimation of crop traits (biophysical variables) from canopy reflectance is the creation of a high-quality training dataset. This can be addressed by using radiative transfer models (RTMs) to generate training dataset representing ‘real-world’ data in situations with varying crop types and growth status as well as various observation configurations. However, this approach can lead to “ill-posed” problems related to assumptions in the sampling strategy and due to uncertainty in the model, resulting in unsatisfactory inversion results for retrieval of target variables. In order to address this problem, this research investigates a practical way to generate higher quality ‘synthetic’ training data by integrating a crop growth model (CGM, in this case APSIM) with an RTM (in this case PROSAIL). This allows control of uncertainties of the RTM by imposing biological constraints on distribution and co-distribution of related variables. Subsequently, the method was theoretically validated on two types of synthetic dataset generated by PROSAIL or the coupling of APSIM and PROSAIL through comparing estimation precision for leaf area index (*LAI*), leaf chlorophyll content (*Cab*), leaf dry matter (*Cm*) and leaf water content (*Cw*). Additionally, the capabilities of current deep learning techniques using high spectral resolution hyperspectral data were investigated. The main findings include: (1) Feedforward neural network (FFNN) provided with appropriate configuration is a promising technique to retrieve crop traits from input features consisting of 1 nm-wide hyperspectral bands across 400-2500 nm range and observation configuration (solar and viewing angles), leading to a precise joint estimation for *LAI* (RMSE=0.061 m^2^ m^-2^), *Cab* (RMSE=1.42 µg cm^-2^), *Cm* (RMSE=0.000176 g cm^-2^) and *Cw* (RMSE=0.000319 g cm^-2^); (2) For the aim of model simplification, a narrower range in 400-1100 nm without observation configuration in input of FFNN model provided less precise estimation for *LAI* (RMSE=0.087 m^2^ m^-2^), *Cab* (RMSE=1.92 µg cm^-2^), *Cm* (RMSE=0.000299 g cm^-2^) and *Cw* (RMSE=0.001271 g cm^-2^); (3) The introduction of biological constraints in training datasets improved FFNN model performance in both average precision and stability, resulting in a much accurate estimation for *LAI* (RMSE=0.006 m^2^ m^-2^), *Cab* (RMSE=0.45 µg cm^-2^), *Cm* (RMSE=0.000039 g cm^-2^) and *Cw* (RMSE=0.000072 g cm^-2^), and this improvement could be further increased by enriching sample diversity in training dataset.

## 1. Introduction

Since the 1960s, using satellite imagery to measure reflectance of surfaces at a scale of tens of metres has been utilised to monitor vegetation health and to attempt to estimate and forecast changes in vegetation cover and condition (Thenkabail et al., 2019). More recently, these imagery methods have been deployed in more proximal sensors (planes, drones, vehicles) that allow analysis of vegetation at higher resolutions (to sub-centimetre scales) in a research field that is sometimes referred to as ‘high-throughput phenotyping’ (HTP) (Chapman et al., 2018). The aim is to indirectly retrieve crop traits such as water and chlorophyll content, with estimation of integrative traits like leaf area index (LAI) gaining the most attention (e.g. Bacour et al., 2002; Jay et al., 2019; Shibayama and Watanabe, 2007; Xu et al., 2019; Yu et al., 2017). HTP methods based on sensor and imaging technologies can rapidly measure a large number of crop traits across time and space in a cost- and labour-efficient way, which can benefit applications in precision agriculture and plant breeding.

Existing retrieval methods can be classified into two major categories depending on where the source of the training data for establishing relationship between target crop trait and spectral signal (canopy reflectance and its derived variables such as vegetation index): (1) statistical methods use observation data collected from practical experiments to build relationship; (2) physical methods either directly use established cause-effect relationship expressed in radiative transfer models (RTMs) or use simulation (synthetic) data generated by these models to rebuild relationship. Compared to statistical methods, the main advantage of physical methods is to allow construction of a training dataset that represents the entire range of possible situations varying in crop types and growth status as well as observation configurations (Baret and Buis, 2008; Dorigo et al., 2007) and consequently provides an easier way to develop more general relationships unrestricted to situations for variable retrieval. There is an increasing interest of the application of ‘model inversion methods’ to RTMs, among of which PROSAIL is the most popular one and has been widely used for variable retrieval (Berger et al., 2018). Within a RTM such as PROSAIL, canopy reflectance across 400-2500 nm range as model output is regulated by input variables including leaf properties, canopy architecture, soil background and observation geometry (viewing and illumination conditions) via radiation absorption and scattering (Jacquemoud et al., 2009). Theoretically, only crop traits presented as input variables in RTMs could be retrieved from model inversion; however, by treating these retrieved variables as intermediate mediums, model inversion can be extended to broader applications, i.e., estimation of leaf properties at canopy level (Campos-Taberner et al., 2018), phenology prediction (J. Xu et al., 2019), land quality evaluation (Wu et al., 2019) and stress detection (Xia et al., 2019).

Model inversion methods are generally subdivided into three sub-categories: numerical optimization approach (e.g., Bacour et al., 2002; Eon et al., 2019; Lunagaria and Patel, 2019), look-up table approach (e.g., Weiss et al., 2000; X. Xu et al., 2019; Zhu et al., 2019), machine learning approach including use of neural networks (e.g., Bacour et al., 2006; García-Haro et al., 2018; Upreti et al., 2019). As summarized in reviews of variable estimation from remote sensing data, different methods have advantages and limitations with no obvious global solution (Baret and Buis, 2008; Dorigo et al., 2007; Verrelst et al., 2015). Although neural networks did not outperform the other approaches for variable retrieval in previous studies (e.g., Combal et al., 2003; Dhakar et al., 2019; Upreti et al., 2019), this may have been due to the low quality of the training dataset without correction using prior knowledge, and/or insufficient utilization of spectral data and/or limitations of the selected neural network algorithms. Compared with other methods, neural networks are theoretically superior in inverting models with massive input variables (such as using hundreds of hyperspectral bands as input to infer canopy variables) and are computationally efficient once the network is fully trained. Thanks to recent developments in deep learning techniques, in this research, we attempted to explore the use of deep learning approach in model inversion for canopy variable retrieval by optimizing network architecture (hyperparameter tuning) and improving training data quality.

Although model inversion methods provide a reasonable way for estimating variables from remote sensing data, none of them can avoid the “ill-posed” problem, namely, the same model output may result from different combination of model input variables. Essentially, the problem is caused by the model uncertainty which results from its simplification of the structure and biochemistry of a canopy, so that more than one state situations of a canopy could result in exactly the same reflectance profile. In practical applications, this problem is aggravated by the poor input parameter selection, i.e. not accounting for bio-physical limitations in the combinations of parameters physically existing in the real-world. However, this problem can be alleviated by using prior knowledge to strengthen constraints on individual variables or between variables. The simplest way is to define the lower and upper values between which the target trait can be retrieved from based on prior information, for example, field measurement data was used for defining input parameter range in the study of Lunagaria and Patel (2019). M. Xu et al. (2019) indirectly introduced constraints between leaf chlorophyll content and LAI by establishing a 2-dimensional matrix-based relationship between leaf chlorophyll content and two vegetation indices (VIs) for VI-based look-up table inversion, which resulted in a better estimation precision than using individual VI. In addition, the utilization of multi-angular observation data in numerical optimization inversion was reported to improve estimation precision of LAI and leaf chlorophyll content (Roosjen et al., 2018) by reducing possible solutions of variable combination via solutions interception.

Linking crop growth model (CGM) to RTM provides a more straightforward solution to address this “ill-posed” problems by directly constraining the sets of RTM input parameters that contribute to canopy reflectance. Theoretically, such an integration method can be implemented in two ways to improve variable retrieval from remote sensing data. The first way is to calibrate/parameterize CGM using canopy reflectance and then using the calibrated CGM to predict target crop traits. Such an application mode can directly retrieve those variables not included in RTMs such as crop yield and also provide variable estimation across the whole growth season (e.g., Guo et al., 2019; Huang et al., 2019; Thorp et al., 2012). The other way is to convert CGM output variables into input variables of RTM and then apply model inversion method on these constrained input variables and corresponding canopy reflectance. This approach has been rarely discussed or explored.

This research focused on estimation of leaf area index (*LAI*), leaf chlorophyll content (*Cab*), leaf dry weight (*Cm*) and leaf water content (*Cw*) of wheat in four locations across Australia wheatbelt. The overall objective was to investigate inversion procedures based on a deep learning approach (feedforward neural network, FFNN) for crop trait estimation, with a special focus on alleviating the “ill-posed” problem in model inversion through linking a CGM (APSIM) and a RTM (PROSAIL) to generate a higher quality training dataset. Firstly, a baseline FFNN model was established to evaluate the use of FFNN for crop trait retrieval. Secondly, this baseline model was used to explore possibility of reduction of hyperspectral bands and effect of observation configuration (solar and viewing angles) for the aim of model simplification. Finally, this simplified model was trained using different datasets generated by PROSAIL or the coupling of APSIM and PROSAIL to investigate the function of model integration by comparing performance of trained FFNNs.

## 2. Methods

### 2.1 Overview

This research contains several key steps as shown in Figure 1. APSIM and PROSAIL models were integrated by passing variables from the former to the latter based on variable transformation relationships. A defined wheat growing space (characterized by genotype, environment and management) and observation conditions (determined by local latitude, day of year and day time) were set up to run APSIM and PROSAIL for simulation of crop traits and canopy reflectance, which resulted in two types of synthetic datasets. The first dataset (PROSAIL dataset) uses the ranges of the input parameters converted from APSIM outputs but allows PROSAIL to be run using samples from full parameter space for any combination of inputs. The second dataset (APSIM-PROSAIL dataset) directly uses input data converted from APSIM outputs to explore a sub-space of input parameters (i.e. limited by the APSIM biology) to run PROSAIL. According to research objectives, these synthetic datasets were reconstructed and used for FFNN training and evaluation in order to explore the possibility of hyperspectral bands reduction, the effect of observation configuration (solar and viewing angles), and the effect of limiting the PROSAIL input parameters to the sub-space as determined by the APSIM.

**Figure 1.**
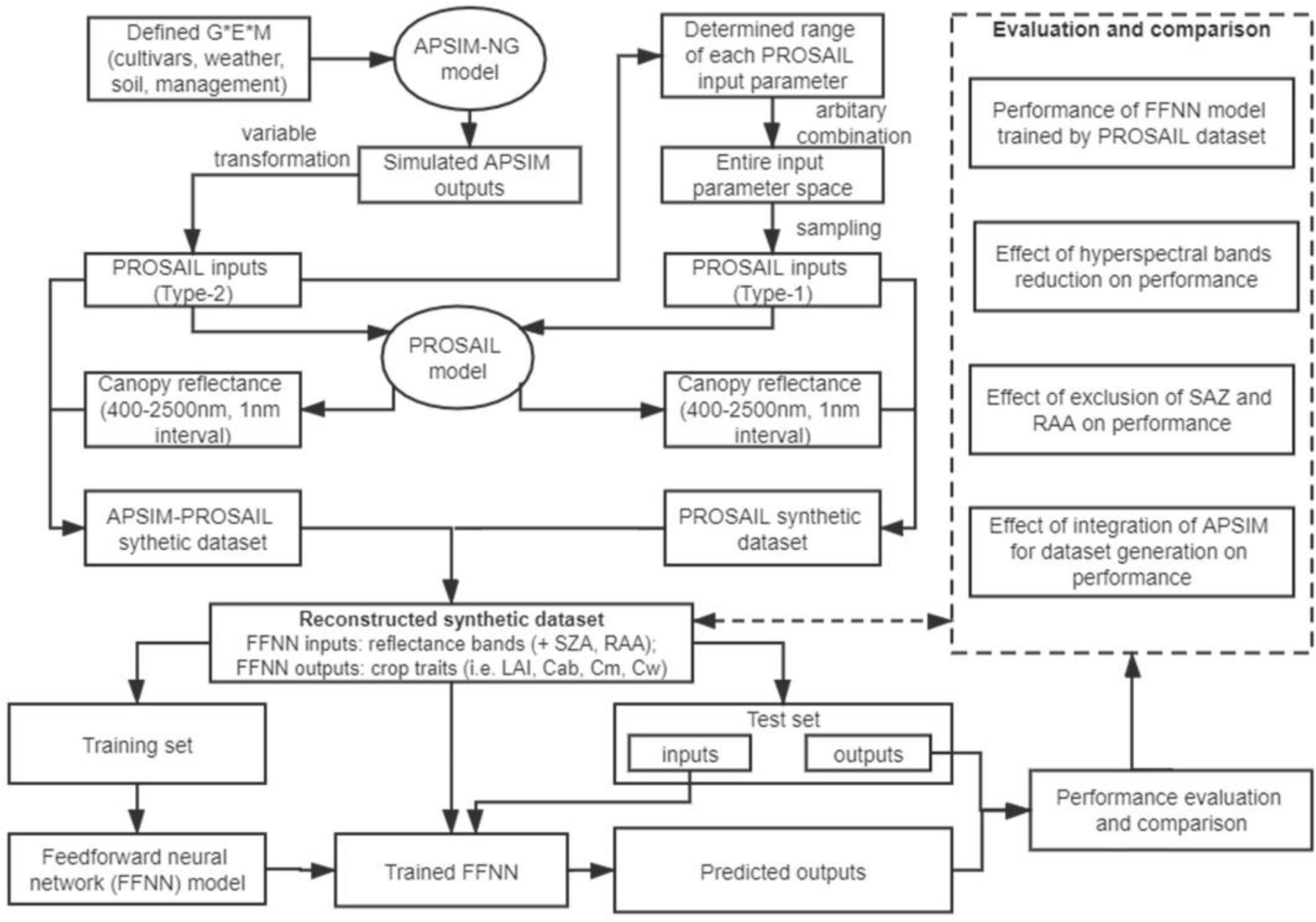
Research flowmap. APSIM-NG denotes Agricultural Production Systems sIMulator (APSIM) Next Generation, which is a crop model. PROSAIL is a radiative transfer model, coupling a leaf optical property model (PROSPECT-D) and a canopy bidirectional reflectance model (4SAIL).

### 2.2 Models and their integration

Agricultural Production System Simulator (APSIM) Next Generation (https://www.apsim.info/apsim-next-generation/) is the new version of APSIM, which is simpler and faster than the classic version (i.e. 7.10., D. Holzworth et al., 2018). APSIM Next Generation or APSIM is driven by major processes in crop physiology and interactions with environment factors and management practices, and widely is used to simulate dynamics of many crop traits (e.g. leaf area index, dry weight of organ parts (i.e. grain, leaf, spike, stem and root)) during growth season at daily scale. APSIM has been validated in many regions around the world (Holzworth et al., 2014).

PROSAIL is the combination of PROSPECT (a leaf optical property model) and SAIL (a canopy bidirectional reflectance model). PROSAIL links the spectral variation of canopy reflectance (mainly related to leaf biochemical contents) with its directional variation (primarily related to canopy architecture and soil/vegetation contrast), which is key to simultaneously estimate canopy biophysical/structural variables such as leaf chlorophyll content and LAI (Jacquemoud et al., 2009). The current version of the PROSAIL model couples PROSPECT-D (see Féret et al., 2017) and 4SAIL (see Berger et al., 2018) and can be downloaded from http://teledetection.ipgp.jussieu.fr/prosail/. Both input and output variables of this PROSAIL model are presented in Table 1. The 14 input parameters can be divided into four categories: leaf properties (*N*, *Cw*, *Cm*, *Cab*, *Car*, *Cant*, *Cbrown*), background soil properties (*rsoil*), canopy architecture (*LAI*, *LIDF*, *hspot*) and solar-object-sensor observation geometry (*SZA*, *VZA*, *RAA*). PROSAIL can output directional canopy reflectance, which is also represented using canopy reflectance or model output without explicit specification in the following sections. For further details about PROSPECT and SAIL model, refer to the original papers (Féret et al., 2017; Jacquemoud and Baret, 1990; Verhoef, 1998, 1984).

**Table 1.**
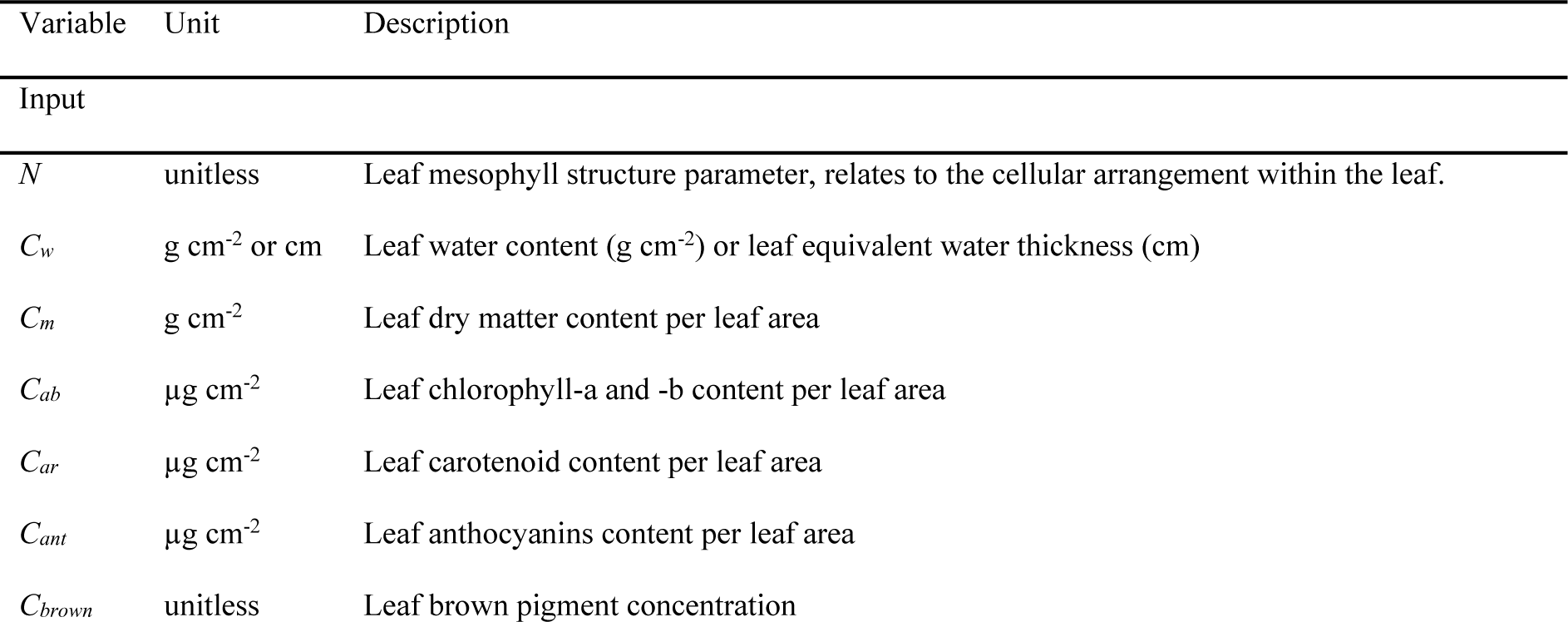
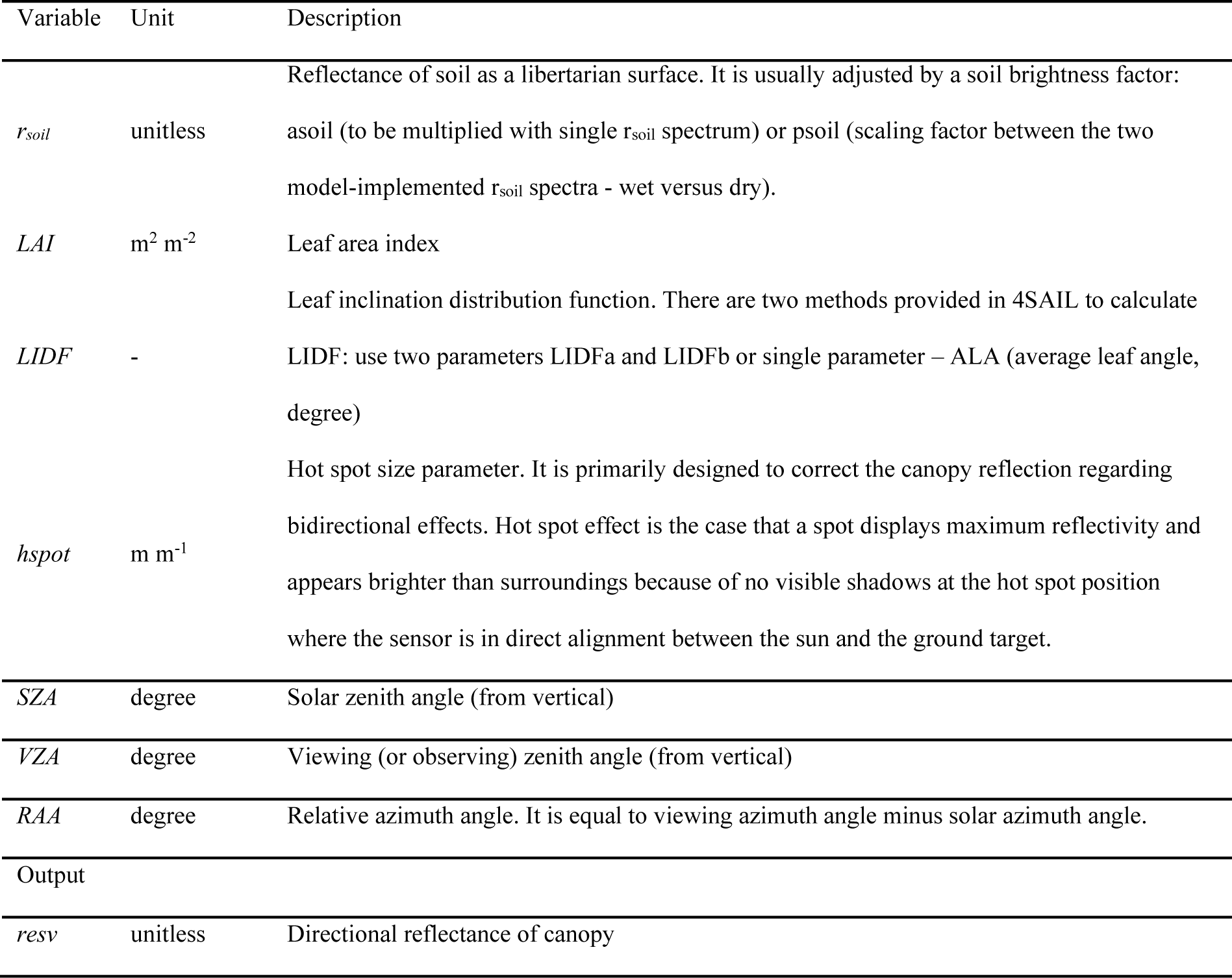
Description of input parameters and output of PROSAIL (PROSPECT-D + 4SAIL)

The coupling of APSIM and PROSAIL is realized by passing output variables of APSIM to PROSAIL as input variables. This permits the coupling model to estimate canopy reflectance from 400 to 2500 nm in 1 nm interval at defined observation conditions (determined by latitude, day of year, and day time) given that required parameters are specified. The transformation of variables is based on a series of equations (Table 2) and more details could be found in Section 1 of supplementary materials.

**Table 2.**
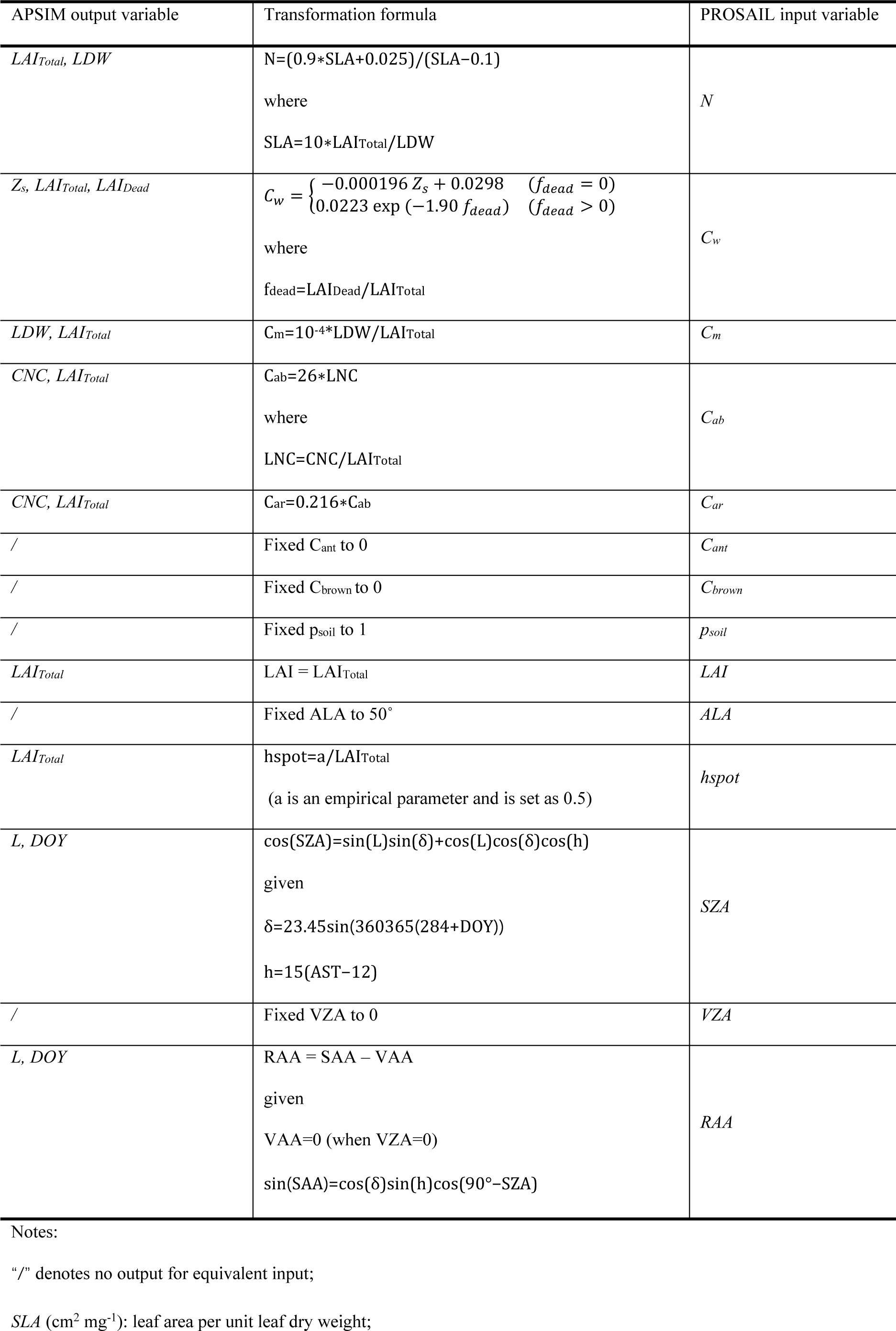
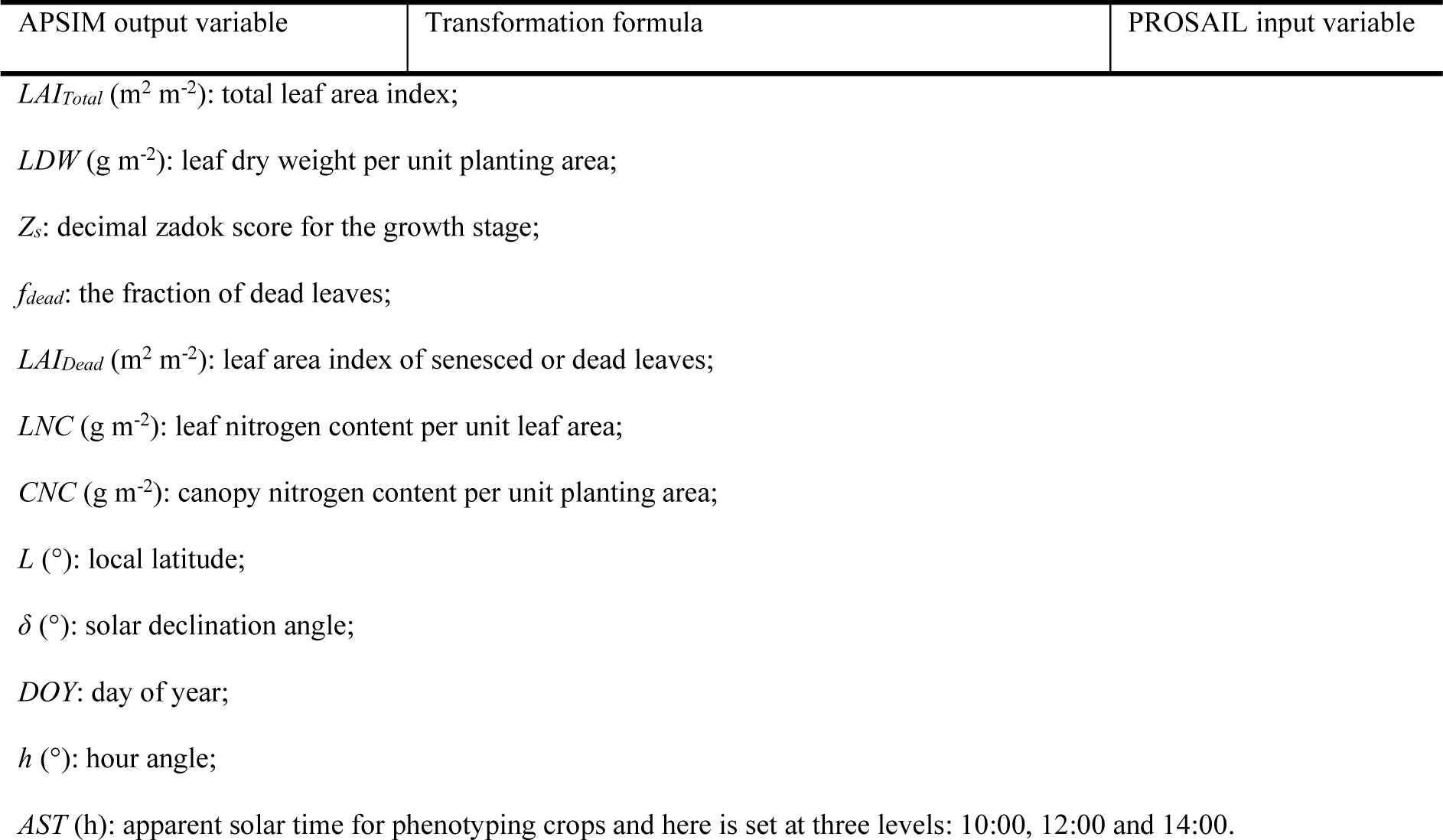
Variable transformation from APSIM output to PROSAIL input.

### 2.3 Synthetic dataset generation

At four sites used to represent diverse conditions across the Australia wheatbelt (Table 3), simulations were run with historical weather records from 2000-2019, the typical soil condition with best initial soil water and local management practices (i.e. fertilization, Table 3, (Chenu et al., 2013)). For each site at each year, 9 cultivars (varying in habit and/or development speed) and 9 sowing dates (from 1-May to 30-June in 1-week interval) were selected to characterize different wheat growth patterns (Table 3). In total, 6 480 simulation seasons were performed using APSIM Next Generation. Major crop traits (Table 2) were output in daily step from emergence to harvest, resulting in 1 080 680 daily records. Based on three defined observation times, these APSIM output records were converted into 3 243 040 PROSAIL input records using variable transformation formulas presented in Table 2, which were then used to determine variation range of each PROSAIL input variable (Table 4).

**Table 3.**
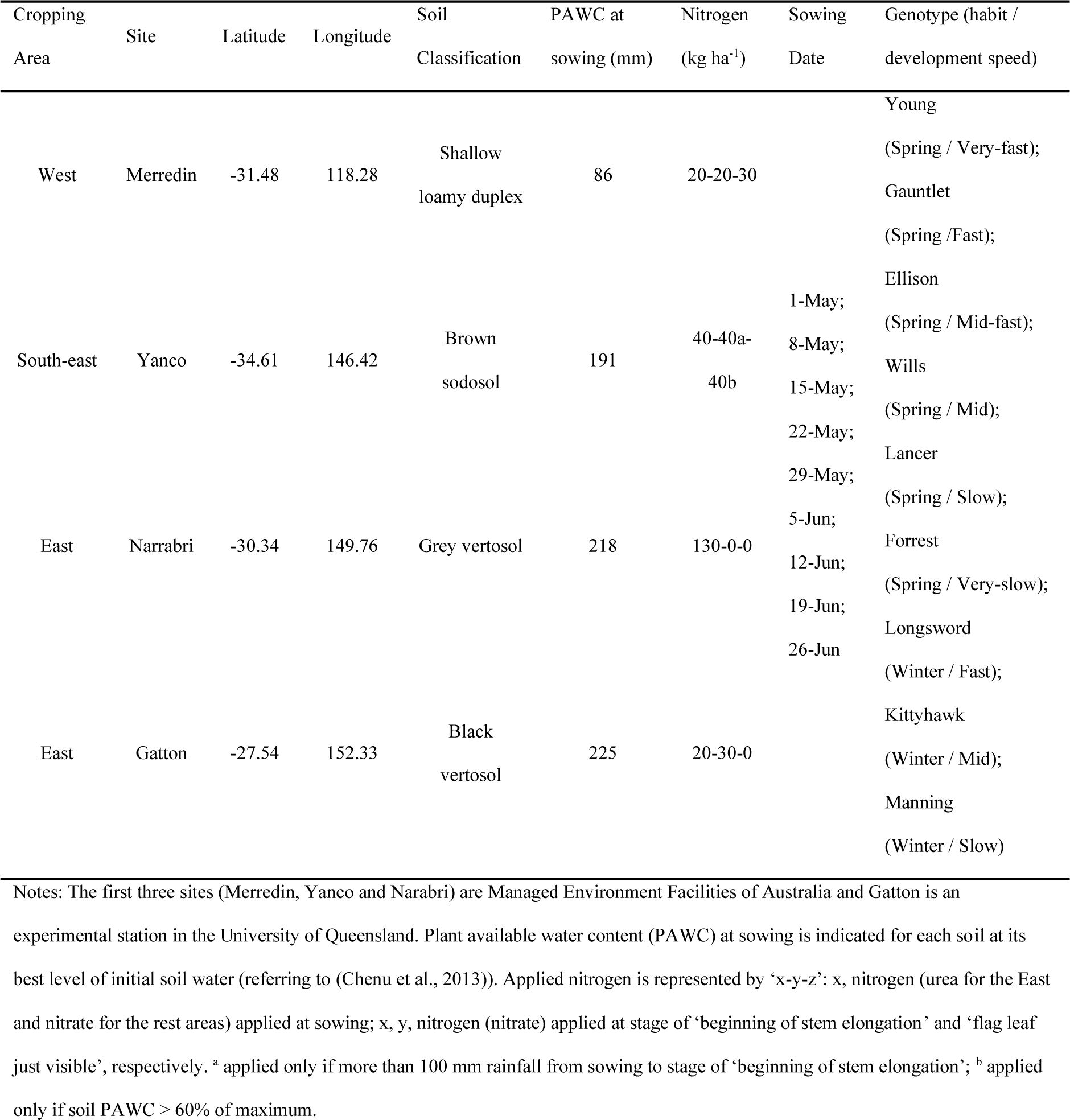
Information of genotype, environment and management used for simulation to represent Australia wheatbelt.

**Table 4.** Setting of PROSAIL input parameters used for sensitivity analysis and synthetic dataset generation.

| Parameters | Local sensitivity analysis (OAT) |  | Global sensitivity analysis (EFAST) |  |  |
| --- | --- | --- | --- | --- | --- |
|  |  |  | / PROSAIL dataset |  |  |
|  | Baseline | Levels | Lower Bound | Upper Bound | Distribution |
| $N$ | 2.25 | 1, 1.6, 2.3, 2.9, 3.5 | 1 | 3.5 | Uniform |
| $C_w$ | 0.017 | 0.003, 0.01, 0.017, 0.023, 0.03 | 0.003 | 0.03 | Uniform |
| $C_m$ | 0.006 | 0.002, 0.004, 0.006, 0.008, 0.01 | 0.002 | 0.01 | Uniform |
|  |  |  | / PROSAIL dataset |  |  |
|  | Baseline | Levels | Lower Bound | Upper Bound | Distribution |
| *C<sub>ab</sub>* | 48 | 5, 26, 48, 79, 90 | 5 | 90 | Uniform |
| *C<sub>ar</sub>* | 10 | 1, 5, 10, 15, 20 | 1 | 20 | Uniform |
| *C<sub>ant</sub>* | 0 | 0 | 0 | 0 | Fixed |
| *C<sub>brown</sub>* | 0 | 0 | 0 | 0 | Fixed |
| *p<sub>soil</sub>* | 1 | 1 | 1 | 1 | Fixed |
| *LAI* | 3.5 | 0.5, 1.5, 2.5, 3.5, 4.5, 5.5, 6.5 | 0.01 | 7 | Uniform |
| *ALA* | 50 | 50 | 50 | 50 | Fixed |
| *hspot* | 1.7 | 0, 0.8, 1.7, 2.6, 3.5 | 0 | 3.5 | uniform |
| *SZA* | 35 | 0, 18, 35, 52, 70 | 0 | 70 | Uniform |
| *VZA* | 0 | 0 | 0 | 0 | Fixed |
| *RAA* | 45 | 0, 22, 45, 68, 90 | 0 | 90 | Uniform |

The synthetic dataset was a set of data pairs constructed from PROSAIL input variables and corresponding outputs. It is an ideal means to use synthetic dataset to demonstrate the application of PROSAIL inversion for estimation of crop traits in theoretical dimensionality, since such a synthetic dataset represents the whole range of possible situations varying in crop types and growth status as well as observation conditions. Based on combination mode of PROSAIL input parameters, the synthetic dataset can be classified into two types: PROSAIL dataset and APSIM-PROSAIL dataset. For PROSAIL dataset, the entire parameter space of input variables consists of arbitrary combination of each parameter changing in their ranges defined in Table 4. The input variables of this type of datasets were generated by sampling a subset from this parameter space based on EFAST’s resampling scheme with given minimum sample size (n). EFAST sensitivity analysis was undertaken multiple times to determine the appropriate n size. Our results show that sensitivity indices can converge to a relative robust value where n ≥ 5000 (Figure S3 in supplementary materials), indicating a subset with sample number of 45000 (=5000*9) is sufficient to reflect relationships between model’s input and output. Compared with input variables of PROSAIL dataset, input variables of APSIM-PROSAIL dataset are not combined arbitrarily but are constrained to wheat growth pattern. The input variables of APSIM-PROSAIL dataset were those directly converted from ASPIM output variables, which contained 2 149 226 unique records after omitting 1 092 814 duplicated ones from the total 3 242 040 records. To our objectives, several synthetic datasets were generated in the way mentioned above (Table 5) and the density distribution and spatial co-distribution of PROSAIL input variables of these datasets in one- or two-dimensional space are presented in Figure 2.

**Figure 2.**
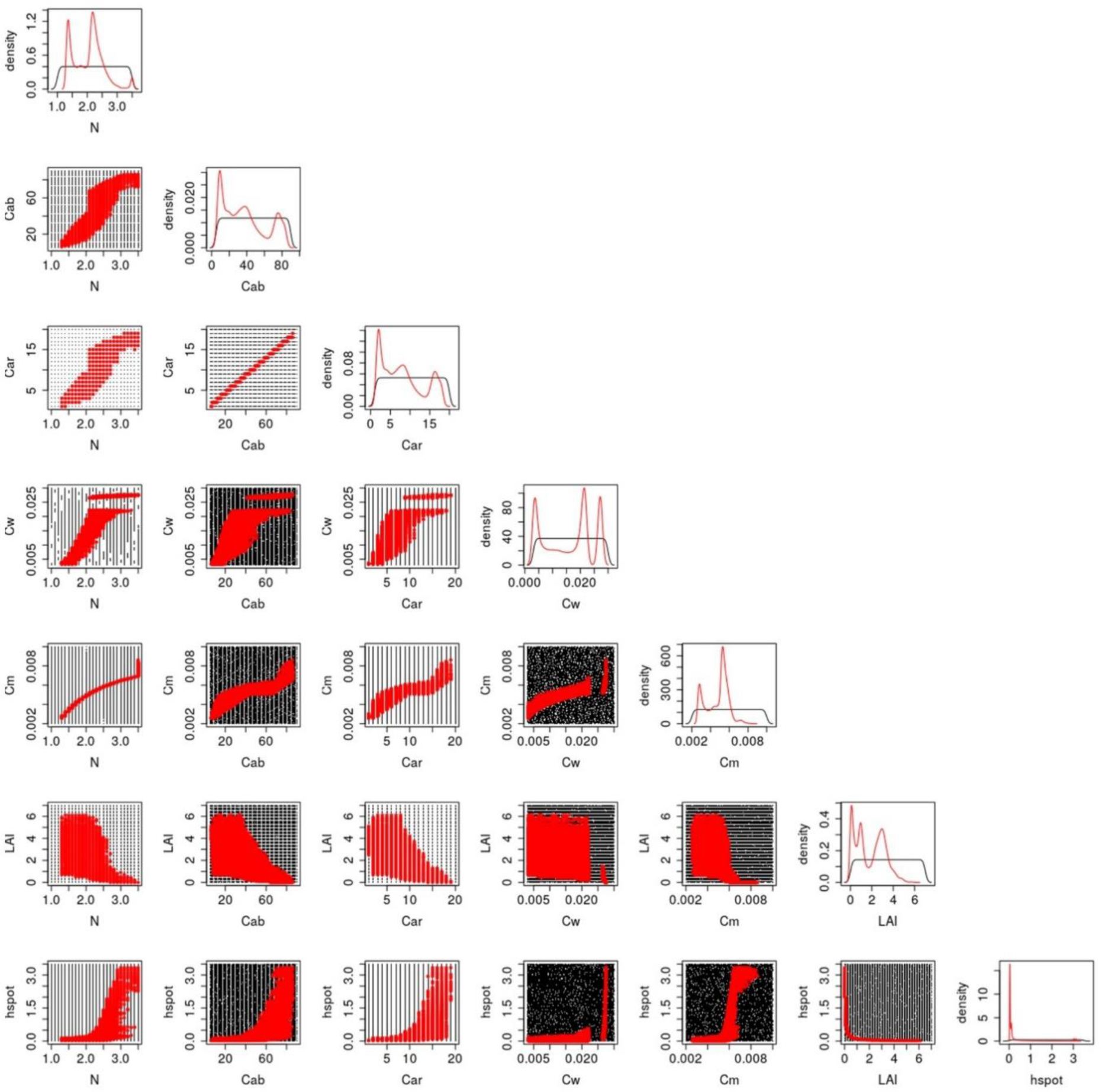
Density distribution and spatial co-distribution of PROSAIL input variables of two types of dataset. Black symbol represents PROSAIL dataset (p_train1) while red symbol represents APSIM-PROSAIL dataset (ap_train1).

**Table 5.** Information of synthetic dataset’s name, type, number and description.

| Dataset<br>Name | Dataset Type | Sample<br>Number | Description |
| --- | --- | --- | --- |
| p_initialTrain | PROSAIL | 45 000 | Sampling from parameter space based on EFAST's resampling scheme with $n=5000$; used to determine FFNN model's hyperparameters in initial training |
| p_test | PROSAIL | 10 000 | Selecting from p_initialTrain; used as test set to evaluate performance of trained FFNN models (trained by p_train1) in experiments SE1-SE7 (see Table 6) |
| p_train1 | PROSAIL | 90 000 | Sampling from parameter space based on EFAST's resampling scheme with $n=10000$; used as train set in in experiments SE1-SE7 (see Table 6) |
| ap_test | APSIM-PROSAIL | 10 000 | Selecting from total 2 149 226 unique samples; used as test set to evaluate performance of trained FFNN models (trained by ap_train1 or ap_train2) in experiment SE8-SE9 (see Table 6) |
| ap_train1 | APSIM-PROSAIL | 90 000 | Selected from the remaining 2 139 226 samples; used as train set in experiment SE8 (see Table 6) |
| ap_train2 | APSIM-PROSAIL | 2 139 226 | The remaining 2 139 226 samples; used as train set in experiment SE9 (see Table 6) |

### 2.4 Sensitivity analysis

For the PROSAIL model, not all canopy reflectance at wavelength from 400 to 2500 nm are sensitive to variation of input parameters. A simplified model with fewer insensitive outputs is superior to the full model for variable retrieval through inversion against a spectral image. For example, inverting variable from a parameter space with fewer possible solutions can improve inversion efficiency and mitigate the ill-posed problem (Verrelst and Rivera, 2017). In order to generate a more representative synthetic dataset used for variable retrieval, a sensitivity analysis was conducted to evaluate the relative importance of each input variable in PROSAIL model and subsequently identify the most influential variables and most sensitive wavelength range.

Sensitivity analysis includes local and global sensitivity analysis. Local sensitivity analysis is commonly referred to “one-at-a-time (OAT)”, which changes one input variable at a time while holding others at their central values for measurement of variation in the model outputs compared with outputs at central point (Verrelst and Rivera, 2017). In contrast, global sensitivity analysis explores the entire variable space and simultaneously changes all variables. The “extended-FAST” (EFAST) method (Saltelli et al., 1999) is a variance-based method and is frequently used in global sensitivity analysis. This method allows the estimation of the first-order and total effect indices for all the input parameters at a total cost of n × p simulations (p is the number of parameters, n is the sample size). First-order effect indices (S_1i_) represent the isolated contribution of i^th^ parameter to the variance of the model output (i.e. canopy reflectance in this study). Total effect indices (S_ti_) represent the total contribution of i^th^ parameter: the isolated contribution of a parameter plus its interactions with other input parameters. The normalized total effect indices (RC_S_ti_) are appropriate to represent the relative contribution of i^th^ input parameter to variation of model output and the normalized first-order effect indices (RC_S_1i_) represent the relative isolated contribution.

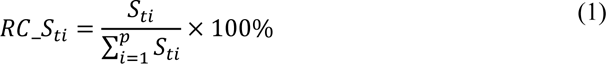

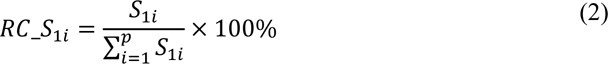

### 2.5 Feedforward neural network (FFNN)

In this research, OAT analysis was undertaken to present how canopy reflectance responds to variation of each input variable in the wavelength range from 400 nm to 2500 nm. In addition, EFAST was chosen as the global sensitivity analysis method to quantify relative contribution of each input variable on canopy reflectance in 400-2500 nm range. Values and ranges of PROSAIL input parameters used for sensitivity analysis were set as shown in Table 4. Feedforward neural networks (FFNNs), also called deep feedforward networks, or multilayer perceptrons, are the quintessential deep learning models (Goodfellow et al., 2016). A FFNN defines a mapping y=f(x;θ) and learns the value of the parameters θ that result in the best function approximation for a prediction: either a classification (discrete) or a regression (continuous). Within this network structure, data passes from the input “x” corresponds to the raw data, which goes through intermediate computations in the function “f” with the parameters “θ”, to the output “y” in a single pass without any feedbacks or cycles.

As presented in Figure 3, FFNN has a multilayer structure consisting of an input layer (the first layer of a network), an output layer (the final layer of a network), and one or more hidden layers (the remaining layers of a network). The total number of layers is called the depth of a network, and each hidden layer of the network consists of many neurons (or units). The dimensionality of these hidden layers determines the width of the network. According to the universal approximation theorem (Cybenkot, 1989; Hornik et al., 1989), a feedforward network can approximate an arbitrary function even with only one hidden layer that is sufficiently wide. However, simply increasing the number of neurons can easily lead to over-parameterization, hence increasing depth seems to be an alternative as experiences from previous studies showed greater depth typically resulted in better generalization (Lin et al., 2014; Zhang et al., 2017). In addition to depth and width, a FFNN has other necessary components: activation, optimizer and loss function. The activation function is a function used to transform data, which allows the layer to learn not only the linear transformation but also the non-linear transformations of the input data and to increase the capacity for better learning of the complex mapping from the input to the output. The optimizer specifies how the training (learning) proceeds through updating model parameters θ (weights) towards a better prediction based on feedback signal from loss function. The magnitude of the move of weights update in training is controlled by learning rate. The loss function quantifies the accuracy of a model on the training data and is used to navigate the training process so as to minimize the training loss. Since a network is trained by iteratively going through the training data, the loss score decreases as training proceeds and finally, it yields a trained network that can accurately estimate the output y with f(x;θ) when consistent minimal loss is observed.

**Figure 3.**
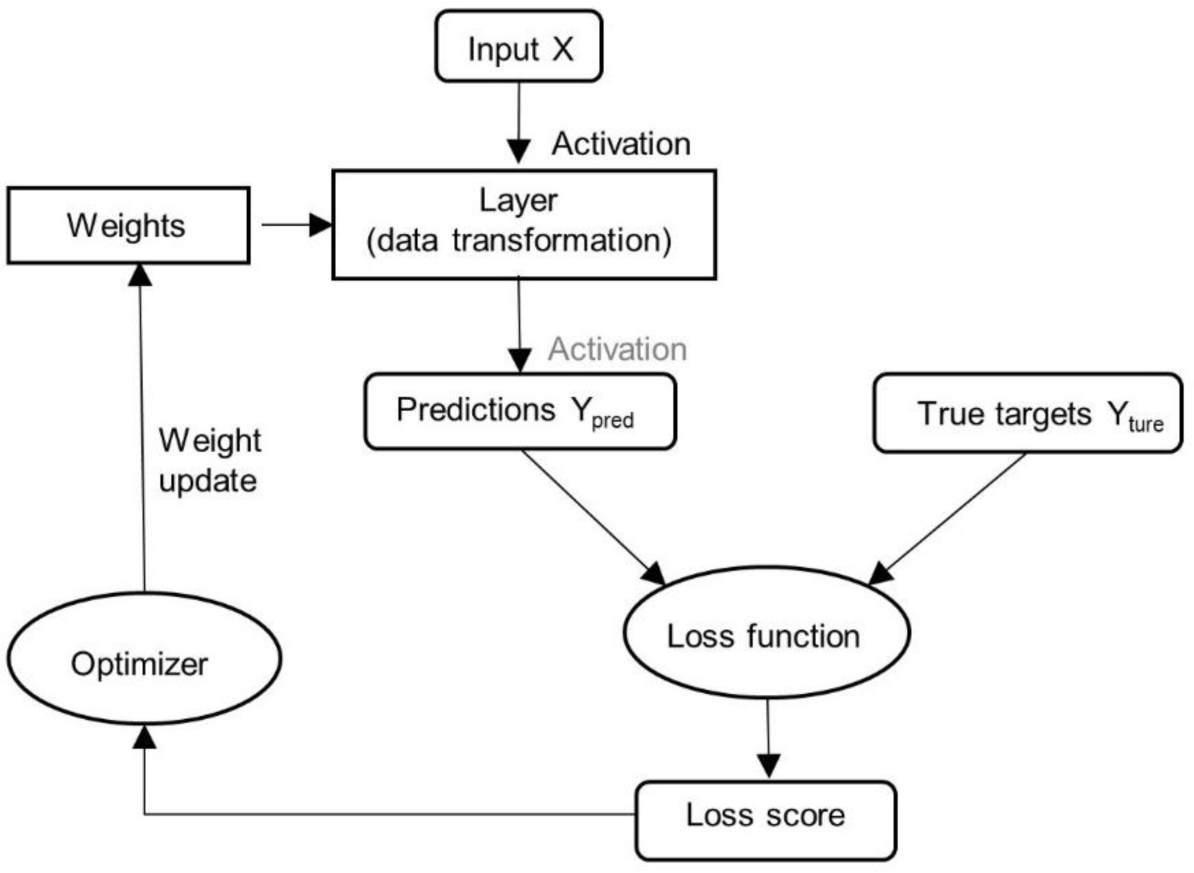
Feedforward neural network working roadmap (adapted from (Chollet, 2017)). ‘Activation’ coloured in black is necessary while that one coloured in grey is unnecessary.

The FFNN model was implemented using Keras in TensorFlow 2.3.0 (https://www.tensorflow.org/). Based on research objectives, the PROSAIL and APSIM-PROSAIL datasets generated in previous steps were reconstructed and several simulation experiments were designed to facilitate other steps in the method (Table 1). In the following sections, we demonstrated how to build, train and evaluate FFNN.

#### 2.5.1 Hyperparameter tuning

Hyperparameters denote variables that govern the training process and the architecture of an FFNN model. Hyperparameters include model hyperparameters (influencing model basic architecture such as the number and width of hidden layers) and algorithm hyperparameters (influencing the speed and quality of training process such as learning rate, activation and optimizer). The process of determining the optimal hyperparameters is called hyperparameter optimization or hyperparameter tuning. For a regression task presented here, the most common loss function is mean squared error (MSE).

In the initial training experiment ‘hypertuning’, dataset ‘p_initialTrain’ (Table 6) was used for hyperparameter tuning, and the best combination of hyperparameters was determined via a two-step optimization process. At the first step, the best combination of learning rate, activation and optimizer was selected by changing these three hyperparameters and holding the number and width of hidden layers to default values (3 hidden layers with 256 units for each layer). At the second step, the best combination of depth and width was chosen based on the algorithm hyperparameters selected at the first step. During this hyperparameter tuning, 3 common values of learning rate, 10 activation functions, 9 optimizers, 5 levels of hidden layer number and 8 levels of unit number were evaluated. In the end, the model structure with 3 hidden layers and 512 units for each hidden layer and using 0.001 as learning rate, ‘softplus’ as activation, ‘Adamax’ as optimizer, was selected as the optimal FFNN structure (Figure S4 and Figure S5) used in later training experiments.

**Table 6.**
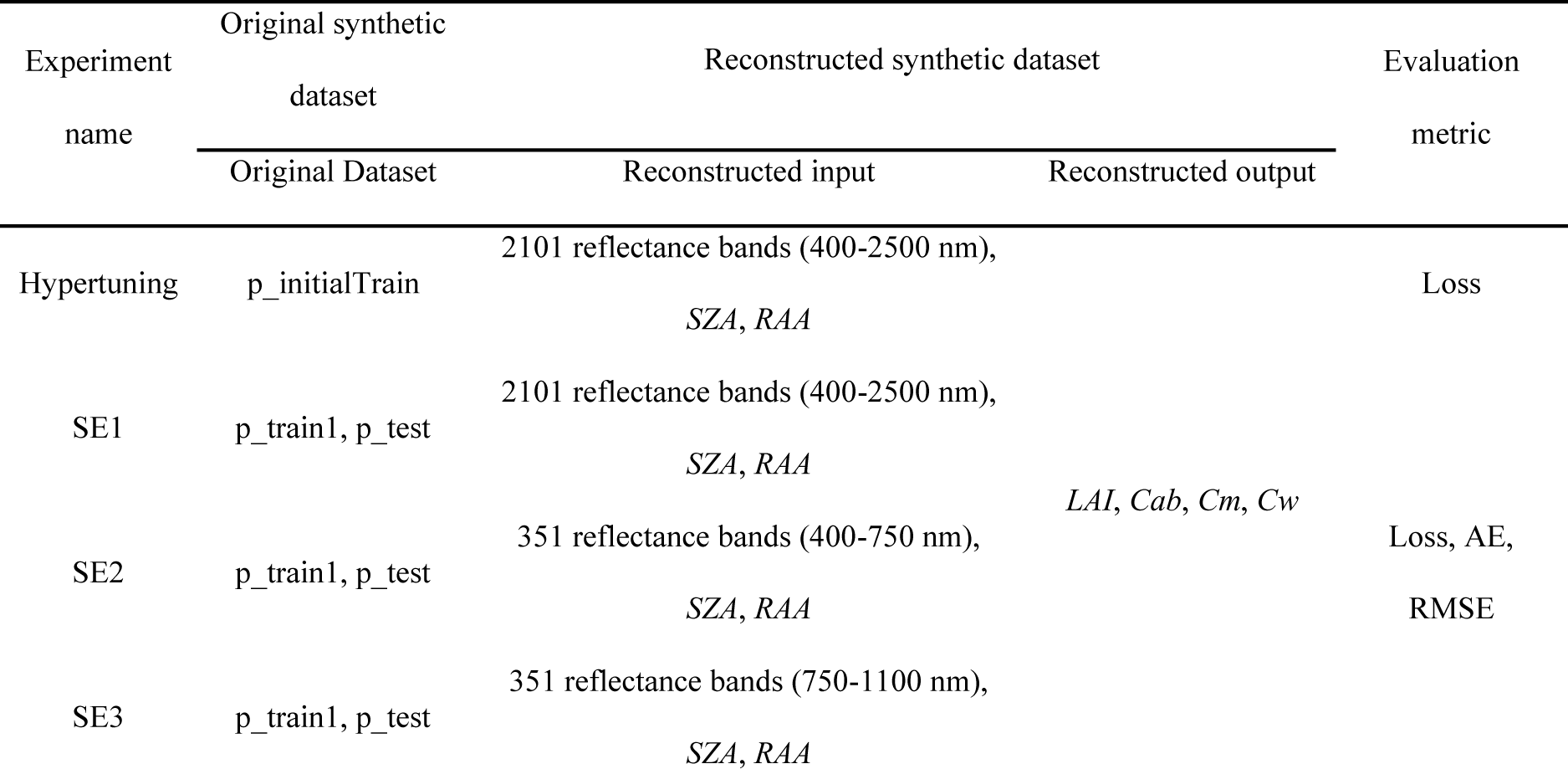
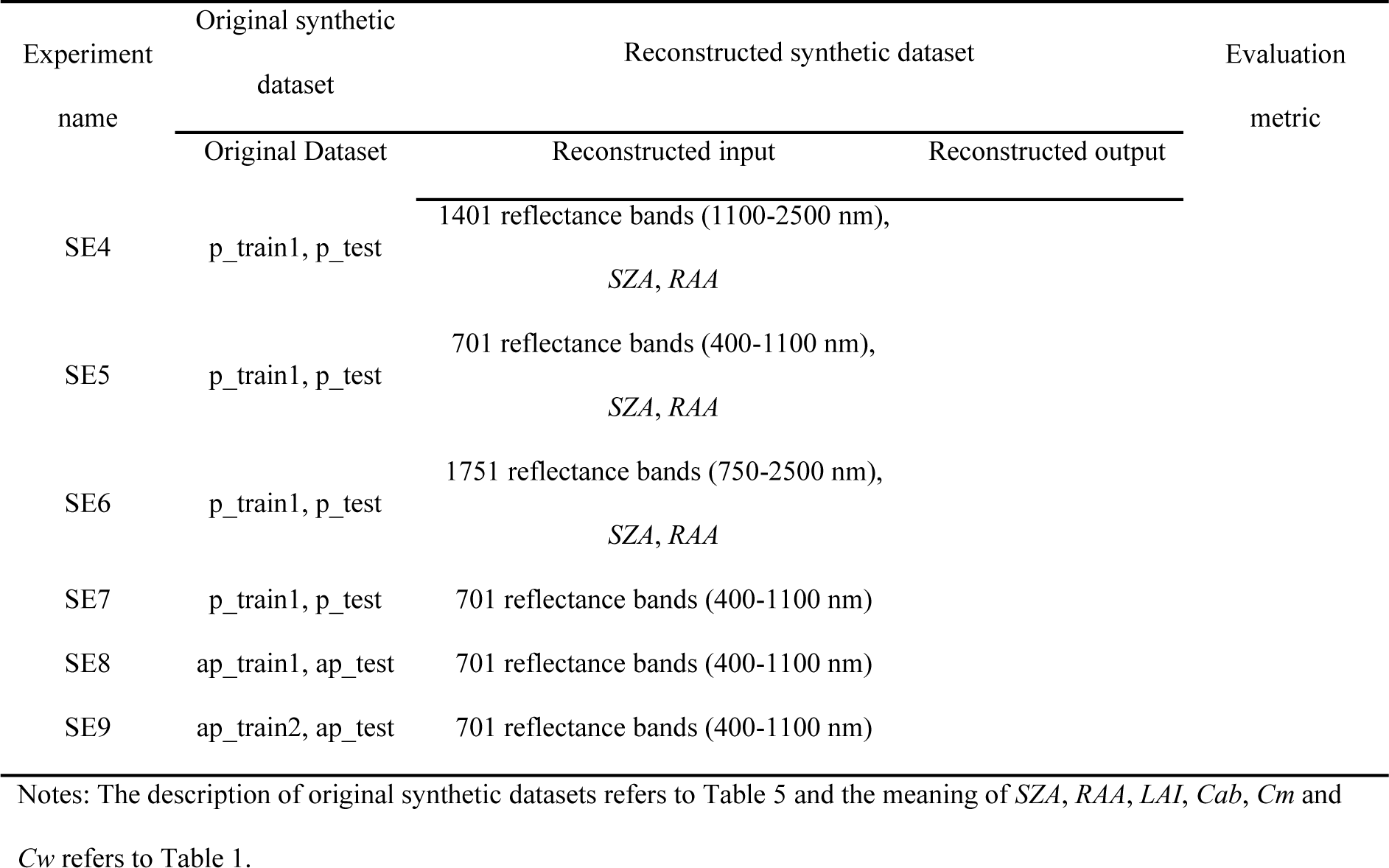
Details of simulation experiment’s name, original and reconstructed synthetic dataset, and evaluation metric.

#### 2.5.2 Training and evaluation

The best model architecture determined in experiment ‘hypertuning’ was used in the following experiments SE1 to SE9. Experiment SE1 was designed to check how well PROSAIL inversion based on FFNN can retrieve target crop traits (i.e. *LAI*, *Cab*, *Cm* and *Cw*) from canopy reflectance. Experiments SE2 to SE6 were used to check whether it is possible to reduce hyperspectral bands used in FFNN’s input on the base of ensuring model’s prediction precision by comparing performance of models using different wavelength range in input. Experiments SE5 and SE7 were used to check whether observation geometry information is necessary to achieve good prediction of crop traits by comparing performance of models including or excluding this information from the input. In order to verify the hypothesis that APSIM-PROSAIL dataset outperforms PROSAIL dataset when being used for traits retrieval, experiments SE7 to SE9 were designed to use different types of datasets for training model.

Loss, the total mean squared error of all output variables after normalization, was used to evaluate FFNN model’s overall performance for joint estimation of all target variables: a smaller loss indicates a higher precision. Absolute error (AE, including its variation range (AE range), standard deviation (std AE) and mean (MAE)) and root mean squared error (RMSE) were used for measurement of each target variable after de-normalization. In particular, AE range and std AE were used to evaluate model’s stability for estimating each target variable while MAE and RMSE were used to evaluate model’s average precision for this variable: a narrower AE range, a smaller std AE, MSE and RMSE indicate a better performance.

## 3. Results and Discussion

### 3.1 Sensitivity analysis of PROSAIL

By applying a local sensitivity analysis (OAT), we characterised a baseline situation for mid-season wheat crop and typical measurement scenario in order to demonstrate the impacts of independently varying input variables on canopy reflectance in 400-2500 nm range (Figure 4). An EFAST analysis qualified relative importance of each variable to total variability of reflectance across 400-2500 nm range in the entire input variable space (Figure 5).

**Figure 4.**
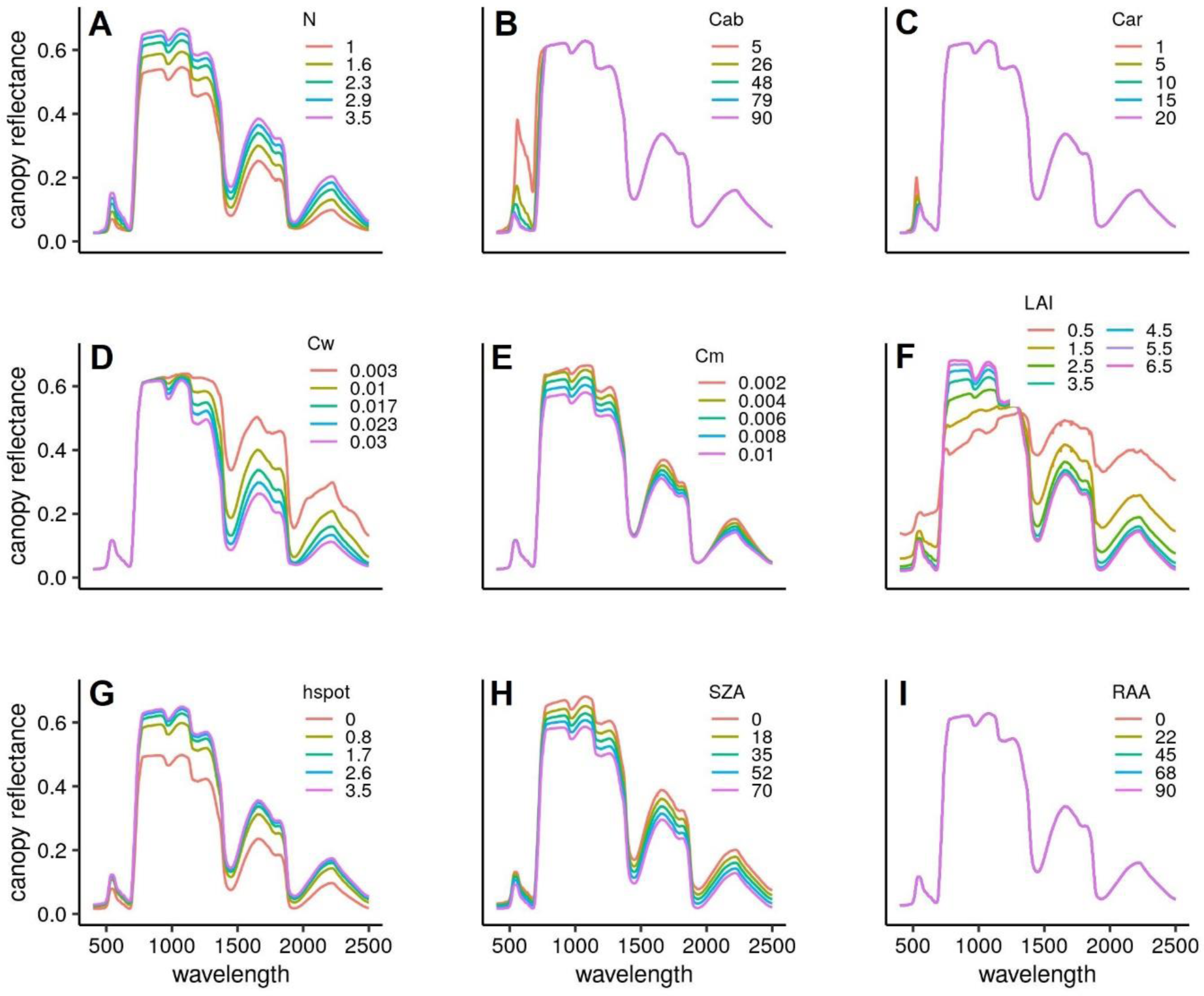
Response of canopy reflectance to change of each parameter. Except for the study parameter changing at different levels, the others are fixed to the baseline according to setting of OAT in Table 3 (Baseline: N=2.25, Cab=48, Car=10, Cant=0, Cbrown=0, psoil=1,Cw=0.017, Cm=0.006, LAI=3.5, ALA=50, hspot=1.7, SZA=35, VZA=0, RAA=45).

**Figure 5.**
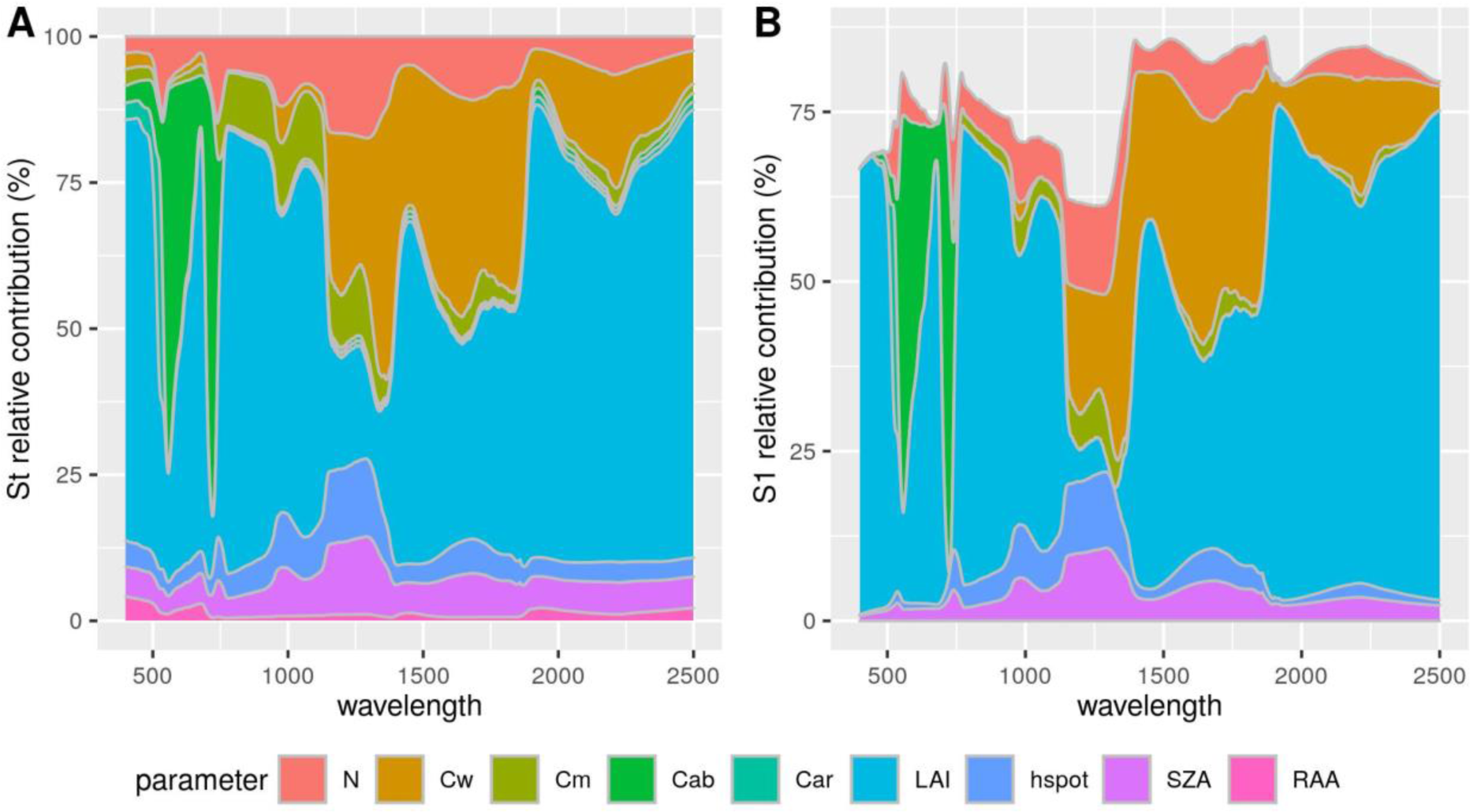
Relative contribution of total effect (A) and first-order effect (B) for each input parameter on canopy reflectance computed by PROSAIL using EFAST global sensitivity analysis

In the OAT, the variables *N*, *hspot*, *Cm* and *SZA* had a consistent either negative or positive effect on reflectance across the whole range (although with small effect of *Cm* in visible region) (Figure 4). Increasing *N* and *hspot* resulted in increases in canopy reflectance (Figure 4A, G) while increasing *Cm* and *SZA* resulted in decreases in canopy reflectance (Figure 4E, H). Increases of two pigment variables *Cab* and *Car* only decreased reflectance in the visible region (Figure 4B, C) where Cab accounted for more than 60% of total variability around 560 nm and 710 nm while influence of *Car* was much less and only in 400-550 nm range (Figure 5). Increasing *Cw* only decreased reflectance in 750-2500 nm range (Figure 4D) and its influence appeared larger at wavelength > 1200 nm (Figure 5), making it become another important driver except for *LAI* in this range. Compared with variables mentioned above, influence of *LAI* was more important and complicated: contributing more than 60% of the total variability especially in range of 400-500 nm, 750-1200 nm as well as 1400-2500 nm (Figure 5) and its increases only increased reflectance in the range around 750-1350 nm but decreased reflectance in the remaining range (Figure 4F). In particular, the contribution of *RAA* was negligible and it only influenced reflectance through interaction with other variables (Figure 5).

Overall, our sensitivity analysis indicated that *LAI* dominated variability of canopy reflectance across 400-2500 nm spectral range, while *Cab* and *Cw* played a key role only at visible range (400-750 nm) and shortwave infrared range (1100-2500 nm), respectively. Similar findings could also be found in other publications (e.g., Danner et al., 2019; Verrelst and Rivera, 2017), although the magnitude of relative contribution of these variables were slightly different due to various method or variables used for sensitivity analysis. The dominance of *LAI* across the whole range is realised via its leaf elements which results in *LAI*. *LAI* indirectly controls the soil reflectance propagating to canopy in low ground cover and controls light absorption in different ranges via leaf optical properties (pigment and water content). Nevertheless, the influence from soil background can be neglected for a canopy with LAI > 3 (Atzberger et al., 2003) in which situation canopy generally reaches a high ground cover (Ramirez-Garcia et al., 2012). The fact that increasing *LAI* decreased total reflectance in 400-750 nm and 1100-2500 nm indicates that photosynthetic pigments (chlorophyll and carotenoid) strongly absorb visible light at wavelengths around 450 nm and 680 nm with leaf water having an influence on light absorption coefficient at wavelength > 1200 nm (Feret et al., 2008).

### 3.2 PROSAIL inversion based on FFNN can predict crop traits from spectral reflectance

To investigate the precision of PROSAIL inversion for variable retrieval, we applied FFNN as model inversion method to retrieve target variables from PROSAIL dataset (p_train1) which was generated by PROSAIL with samples from full parameter space for arbitrary combination of input variables (Figure 1). Performance of the trained FFNN model for target variable retrieval from canopy reflectance in 400-2500 nm range and observation geometry information (here is *SZA* and *RAA*) is reported in Table 7 (results of SE1), Figure 6 and Figure 7. The small values of MAE and RMSE show the trained FFNN reached high precision for joint estimation of *LAI*, *Cab*, *Cm* and *Cab* when the input variables were allowed to explore all combinations across valid physiological ranges. However, the uncertainty range of estimations were occasionally large, for instance, the absolute error of 95% samples for *Cab* estimation was within 2.19 µg cm^-2^ while the maximum error was more than 10 times bigger up to 26.92 µg cm^-2^ (see SE1 in Table 7). The same issue occurred to estimation for the other three variables. Additionally, the trained FFNN continued to make good estimations at different levels of the true value, although larger true values tended to have a higher probability of larger absolute errors (Figure 6).

**Figure 6.**
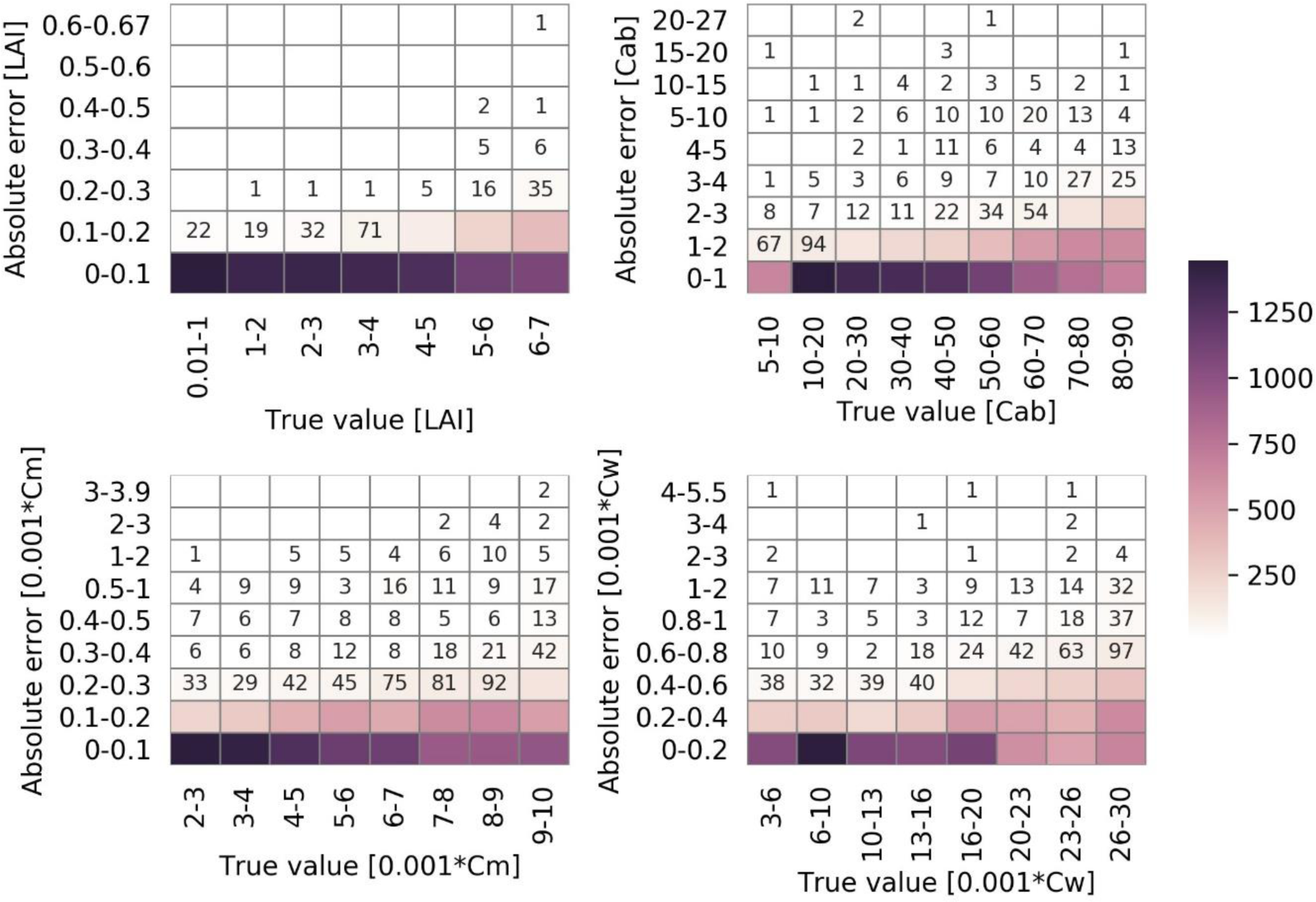
Heatmap of model performance at different levels (SE1). Numbers in cells of the figure show the number of samples whose prediction absolute errors of true values in a range were within a specific range. Numbers less than 100 are shown to differ small values from zero. The unit of absolute error (true value) of LAI, Cab, Cm and Cw is m^2^ m^-2^, µg cm^-2^, g cm^-2^ and g cm^-2^, respectively.

**Figure 7.**
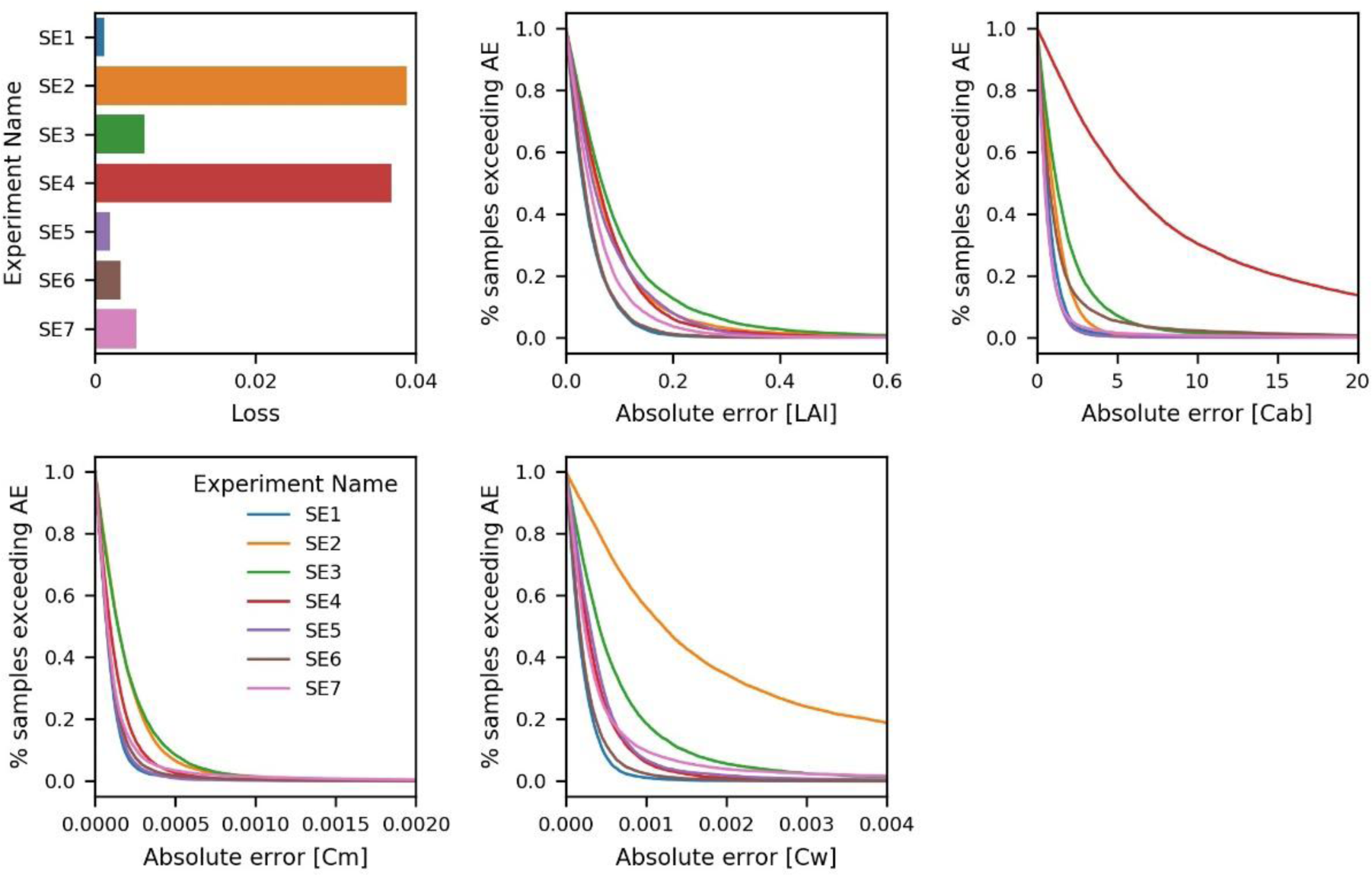
Total performance and its empirical distribution for models with different input features. Loss is a unitless indicator and represents the total mean squared error of joint estimation of four model output variables after normalization. Absolute error (AE) is the difference between the true value of each variable and its prediction after de-normalization. The unit of AE of LAI, Cab, Cm and Cw is m^2^ m^-2^, µg cm^-2^, g cm^-2^ and g cm^-2^, respectively.

**Table 7.**
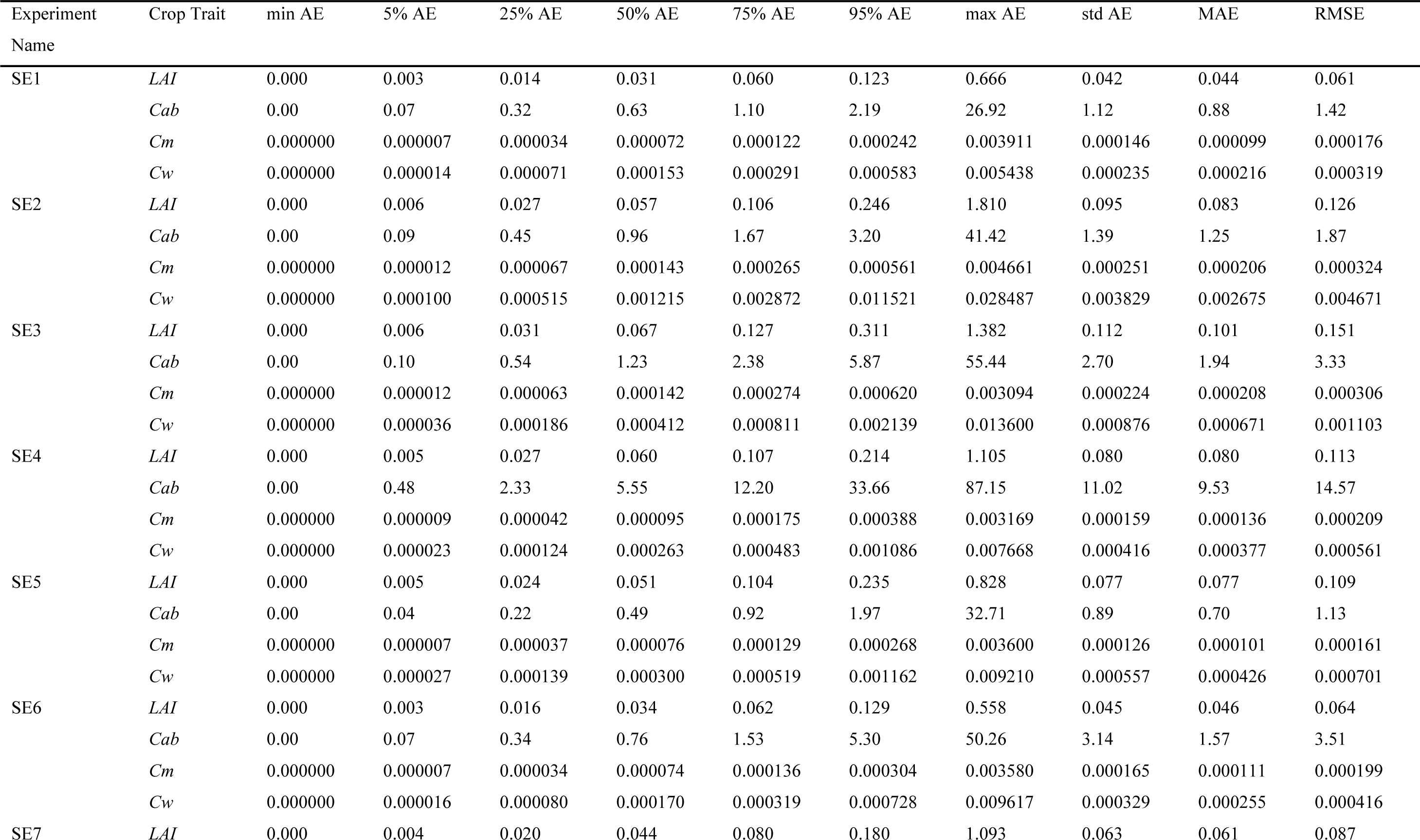
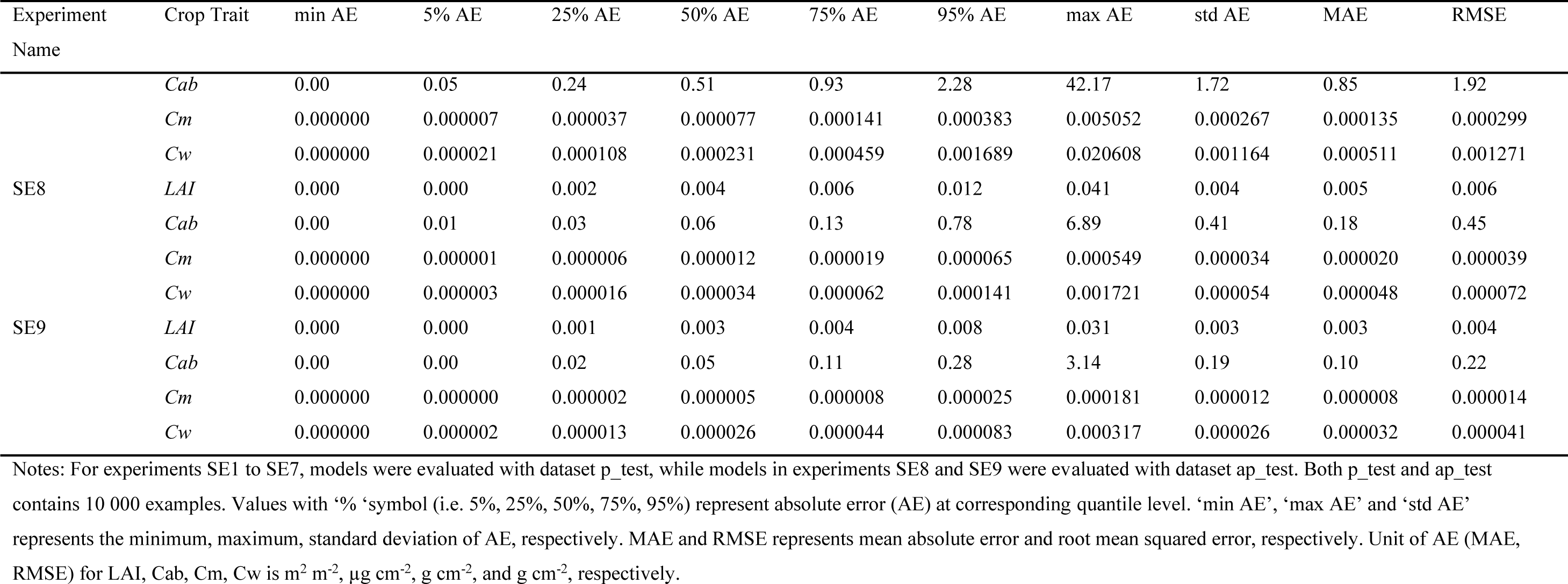
Statistical information of model’s prediction error for each variable in each experiment.

The trained model might not perform as well as presented here when it was applied to retrieve variables from real observation data due to measurement and model uncertainties. Thus, it would be unfair to compare our results with those from real observation data and should be more reasonable to compare with those also from simulation data. As presented in Table 8, our results were favourable, with approximately 10 times smaller RMSE for estimation of *LAI*, *Cab*, *Cm* and *Cw*, compared with simulated results from other model inversion studies. Reasons for such a good estimation exhibited in our research are likely due to the use of better architecture and algorithm used in neural network as well as more complete information included in massive hyperspectral bands as other studies inverted variables from only a few broad/narrow bands (Atzberger, 2004; Baret et al., 2007; Upreti et al., 2019) or derived VIs (le Maire et al., 2008; M. Xu et al., 2019). This result highlights the advantages of new deep learning techniques applied to high spectral resolution hyperspectral data and, demonstrate that it is viable to use deep learning approach to invert hyperspectral data to retrieve variables.

**Table 8.** Root mean squared error (RMSE) of target variable estimation from this and other inversion studies.

| LAI ($m^2 m^{-2}$) | Cab ($\mu g cm^{-2}$) | Cm ($g cm^{-2}$) | Cw ($g cm^{-2}$) | Inversion method | Reference |
| --- | --- | --- | --- | --- | --- |
| 0.81 (0.59) | 9.94 (7.84) | / | 0.0037 (0.0028) | Neural network | Atzberger, 2004 |
| 1.10 | / | / | / | Neural network | Baret et al., 2007 |
| 1.31 | 9.84 | 0.001414 | / | VI empirical regression | le Maire et al., 2008 |
| / | 10.5 (7.9) | / | / | VI-based look-up table | M. Xu et al., 2019 |
| 0.99 | 11.4 | / | / | Bagging trees |  |
| 1.04 | 11.02 | / | / | Neural network in ARTMO |  |
| 1 | 11.55 | / | / | Random forest tree bagger | Upreti et al., 2019 |
| 1.18 | 11.16 | / | / | Partial least square regression |  |

| $LAI$ ($m^2 m^{-2}$) | $Cab$<br>($\mu g cm^{-2}$) | $Cm$ ($g cm^{-2}$) | $Cw$ ($g cm^{-2}$) | Inversion method | Reference |
| --- | --- | --- | --- | --- | --- |
| 1.18 | 11.15 | / | / | Least square linear regression |  |
| 1.14 | 11.61 | / | / | Boosting trees |  |
| 1.3 | 14.99 | / | / | Regression trees |  |
| 1.4 | 16.03 | / | / | Random forest fit ensemble |  |
| 0.94 | 10.55 | / | / | Gaussian process regression |  |
| 0.061 | 1.42 | 0.000176 | 0.000319 | Feedforward neural network | SE1 in Table 7 |
| 0.087 | 1.92 | 0.000299 | 0.001271 |  | SE7 in Table 7 |

Notes: Figures in brackets are the RMSEs after improvement by using a more novel estimation approach. In Atzberger's study, an object-based method was introduced to add spatial constraints on variables. In the study of M. Xu et al, a matrix-based method was used to constrain co-distribution of $LAI$ and $Cab$ .

### 3.3 FFNN could still achieved compatible prediction after reducing wavelength bands and excluding observation geometry in model input

We have demonstrated trained FFNN can well predict crop traits from input features consisting of spectral reflectance in 400-2500 nm range and observation configuration (solar and viewing angles) and in this section, we explored the possibility of hyperspectral bands reduction and effect of observation configuration in order to simplify FFNN model (Figure 1).

There are two main reasons to investigate effects of the use of reflectance bands in different ranges for variable retrieval. Firstly, most sensors used currently only cover part of the whole 400 to 2500 nm wavelength range. Secondly, a large number of reflectance bands used as input to FFNN model results in a complicated model structure, which might make it less suitable for practical application. Considering both the wavelength range classification and ranges covered in current hyperspectral sensors, the model was retrained using five sets of wavelength ranges: 400-750 nm, 750-1100 nm, 1100-2500 nm, 400-1100 nm, and 750-2500 nm. Performance of trained FFNN using reflectance bands in these ranges alone are shown in Table 6 (results of SE 2 to S6) and Figure 7. It is clear that reflectance bands in 400-750 nm and 1100-2500 nm were not suitable for joint estimation of target variables (Figure 7). That is because the use of reflectance bands in these two ranges produced poor estimation for *Cab* and *Cw*, respectively, in which ranges canopy reflectance was insensitive to variation of *Cab* or *Cw* as demonstrated in Section 3.1. For joint retrieval of four target variables, the range of 400-1100 nm could be used as an alternative to replace the whole range for the aim of model simplification: the use of reflectance bands in this narrower range provided nearly same good estimation for *Cab* and *Cm* but less precise estimation for *LAI* and *Cw* by comparing SE1 and SE5.

The reason to attempt to exclude observation configuration (solar and viewing angles) from input of FFNN model is based on two considerations. One consideration is about information necessity and redundancy: the influence of observation configuration on canopy reflectance is likely to be implicitly reflected in hundreds of reflectance bands, so the observation configuration is unnecessary if a large number of reflectance bands are used in FFNN inputs. The other consideration is about information acquisition and availability: observation condition (solar and viewing angles) varies across pixels for an image, so observation configuration of each pixel must be calculated alone from this pixel’s latitude, DOY and day time for phenotyping this pixel if precise retrieval using this method is expected at pixel-level. However, the gaining of observation configuration for every pixel is normally inaccessible especially for an image with large cover consisting of massive number of pixels. By comparing results of SE5 (400-1100 nm with solar/viewing angles) and SE7 (400-1100 nm without angles), it shows that exclusion of observation configuration in input of FFNN model just slightly decreased estimation precision for retrieval of *Cab*, *Cm* and *Cw* and even slightly increased overall estimation for *LAI* (RMSE was 0.109 µg cm^-2^ for SE5 and 0.087 µg cm^-2^ for SE7). Regardless, the results show that the trained FFNN model with input only including reflectance bands in 400-1100 nm range without angles still produced much smaller RMSE than those in previous model inversion studies for variable retrieval (see Table 8).

### 3.4 Coupling APSIM and PROSAIL can further improve FFNN’s prediction precision

While the FFNN model was able to perform well when trained across sets of variables that explore the entire ranges of realistic values, the key part of this paper was to investigate the impact of constraining the input variables to be limited to ‘physiologically realistic’ combinations (Figure 1). Figure 8 indicates that the use of APSIM-PROSAIL dataset (ap_train1 and ap_train2) generated by coupling APSIM and PROSAIL significantly improved FFNN’s performance for estimation of all target variables using reflectance bands in 400-1100 nm without observation configuration (solar/viewing angles). This improvement of model performance presented in both average precision (smaller MAE and RMSE) and stability (narrower uncertainty range, smaller standard deviation of AE) according to statistical results in Table 7. For instance, compared with results using PROSAIL dataset (p_train1, SE7), the use of APSIM-PROSAIL dataset (ap_train1, SE8) for *LAI* estimation narrowed the uncertainty range of AE from 0∼1.093 m^2^ m^-2^ to 0∼0.041 m^2^ m^-2^ and also reduced standard deviation of AE (from 0.063 to 0.004 m^2^ m^-2^), MAE (from 0.061 to 0.005 m^2^ m^-2^) and RMSE (from 0.087 to 0.006 m^2^ m^-2^). The magnitude of improvement resulting from model integration is much larger than that resulting from adding spatial constraints on related variables as demonstrated in the study of Atzberger (2004) where RMSE reduced from 0.81 to 0.59 m^2^ m^-2^ for LAI estimation as exhibited in Table 8. It deserves to emphasize that this improvement is irrelevant to the number of samples used for training (sample size of p_train1 and ap_train1 is the same) and increasing sample size of training set tended to improve model performance in further (sample size of ap_train2 (2 139 226) is larger than ap_train1 (90 000)).

**Figure 8.**
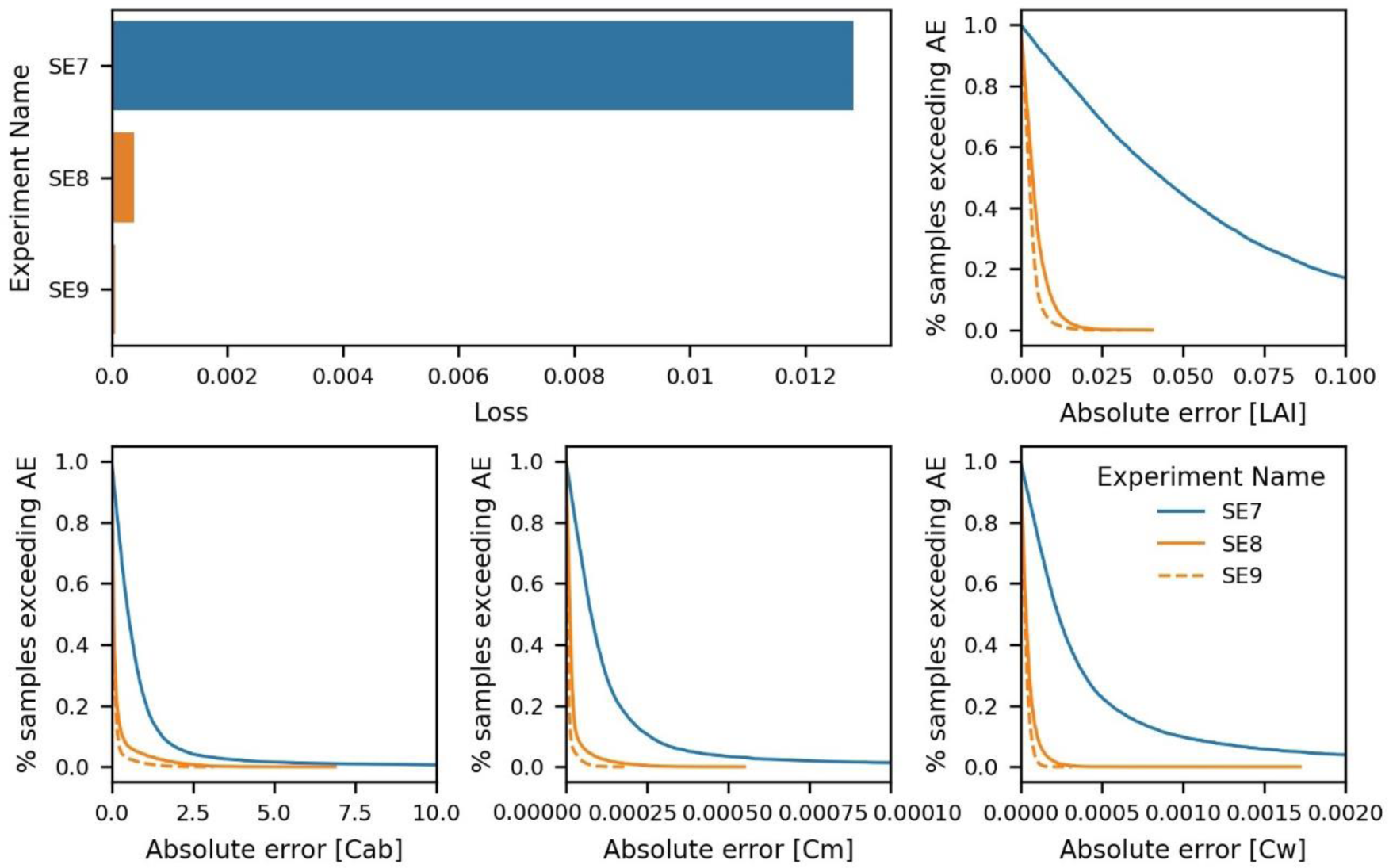
Total performance and its empirical distribution for models trained using different datasets. Loss is a unitless indicator and represents the total mean squared error of joint estimation of four model output variables after normalization. Absolute error (AE) is the difference between the true value of each variable and its prediction after de-normalization. The unit of AE of LAI, Cab, Cm and Cw is m^2^ m^-2^, µg cm^-2^, g cm^-2^ and g cm^-2^, respectively.

The success of this integration in building improved inverse models seems to lie in providing a higher quality training dataset as FFNN totally learns from data provided. CGMs are powerful to predict crop growth and development characterised by a series of physiological processes, thus using CGM to produce sets of input variables of RTM can create a distribution and co-distribution of related variables closer to real situations defining canopy architecture and potentially remove cases that can be simulated but may not be actually observed (sampled) in a real-world from training dataset. Figure 2 clearly shows difference of variable distribution in two types of training dataset whose input parameters vary in the same range. It should be noticed that the difference of distributions projected in low-dimensional space of two datasets can indicate that their distributions projected in high-dimensional space must be different, but that a similarity between distributions in low-dimensional space does not guarantee that their distributions in high-dimensional space are also the same.

The continuous improvement in model performance as sample size increased, it is likely associated to a richer diversity of samples rather than growing number of same samples. Adding more existing samples in training dataset does not enhance learning of relationships between variables within a network. It is reported that a uniform distribution of the variables may obtain a more even distribution of the uncertainties in spite of a poor RMSE (Baret and Buis, 2008). However, our results show a non-uniform distribution closer to true distribution (variables in APSIM-PROSAIL dataset) can also obtain a relative even uncertainty distribution (see Figure S6 and Figure S7) and a better RMSE (see SE8 and SE9 in Table 8) compared with results from PROSAIL dataset with uniform distribution variables (see Figure 6 and SE7 in Table 8). A small consequence is that there is a higher probability of larger error when estimating larger true value. Overall, these findings offered evidence to support that integrating CGM and RTM is a promising way for variable retrieval from hyperspectral data in theoretical dimensionality.

## 4. Conclusion

In general, a completed workflow of integrating APSIM and PROSAIL was demonstrated in this research, resulting in an applicable coupling APSIM-PROSAIL model. Additionally, this research also illustrated the process of choosing appropriate hyperparameter for FFNN model and presented the advantages of using FFNN for crop traits retrieval. It was also demonstrated that the model could be applied to subsets of different wavelength ranges in order to suit different types of instrumentation and applications. However, the major contribution of this research is to demonstrate a practical way to generate higher quality training data which can better characterise the real canopy realization and, prove its practicability in theory. Although this trained FFNN model might not perform as well as presented here when it is applied to retrieve variables from real observation data due to measurement and model uncertainties, it is expected to be able to make relative good performance according to difference of estimation precision on simulated and observed data from other model inversion studies. In future, we are going to investigate the performance of trained FFNN model using simulation data in different simulated situations as well as practical production environments.

## Author’s responsibility

QC implemented this research and wrote the draft of the manuscript. SC helped formulate research ideas and BZ provided guidance of use of APSIM Next Generation model and outlined the framework of this paper with QC. TC provided useful technical advices in the application of the deep learning models. All authors engaged in reviewing and editing this manuscript.

## Acknowledgement

QC is receiving a scholarship co-funded by the China Scholarship Council and the University of Queensland and has been supported by CSIRO as a visiting student.

