## Supplementary_materials for "Integration of APSIM and PROSAIL models to develop more precise radiometric estimation of crop traits using deep learning" (from another version of this preprint: version 1, 2021-02-03)

### Section 1 APSIM-PROSAIL coupling model

The APSIM-PROSAIL coupling model is established by passing output variables of APSIM to PROSAIL as input variables. This permits APSIM-PROSAIL to estimate canopy reflectance from 400 to 2500 nm in 1 nm interval on daily step given that required parameters are specified. The transformation of variables is based on a series of equations below.

The leaf structure parameter (N) can be calculated from specific leaf area (SLA, $cm^2$ $mg^{-1}$) according to the description in (Jacquemoud and Baret, 1990).

$$N = (0.9 * SLA + 0.025)/(SLA - 0.1) \quad (1)$$

$$SLA = 10 * \mathrm{LAI}_{Total}/LDW \quad (2)$$

where $\mathrm{LAI}_{\mathrm{Total}}$ ($m^2$ $m^{-2}$) represents the total leaf area index; LDW (g $m^{-2}$) represents the leaf dry weight per unit planting area.

The leaf dry matter content per leaf area ($C_m$, g $cm^{-2}$) can be calculated from LDW and $\mathrm{LAI}_{\mathrm{Total}}$, as shown in Eq (3).

$$C_m = 10^{-4} * LDW/\mathrm{LAI}_{Total} \quad (3)$$

Previous studies have reported there is a strong linear relationship between leaf nitrogen content (LNC, g $m^{-2}$), leaf chlorophyll content ($C_{ab}$, μg $cm^{-2}$) and leaf carotenoid content ($C_{ar}$, μg $cm^{-2}$) (Table S1).

*Table S1 Relationship between leaf nitrogen, chlorophyll and carotenoid content*

| Relationship between $C_{ab}$ (μg $cm^{-2}$) and LNC (g $m^{-2}$) | Relationship between $C_{ar}$ (μg $cm^{-2}$) and $C_{ab}$ (μg $m^{-2}$) | Plant types | References |
|---|---|---|---|
| $C_{ab} = 30.75^a * LNC$ | $C_{ar} = 0.2159^c * C_{ab}$ | Woody and herbaceous species | (Jacquemoud et al., 1996) |
| $C_{ab} = 26.00^b * LNC$ | $C_{ar} = 0.2160^c * C_{ab}$ | Wheat | (Yang et al., 2015) |

| Relationship between $C_{ab}$ ($\mu g\ cm^{-2}$) and LNC ($g\ m^{-2}$) | Relationship between $C_{ar}$ ($\mu g\ cm^{-2}$) and $C_{ab}$ ($\mu g\ m^{-2}$) | Plant types | References |
|---|---|---|---|

[a] the ratio of average $C_{ab}$ and average LNC;

[b] the regression coefficient of $C_{ab}$ and LNC;

[c] the ratio of average $C_{ar}$ and average $C_{ab}$.

Based on Table S1, for wheat, $C_{ab}$ and $C_{ar}$ can be calculated using Eq (4) and Eq (5), respectively.

$$C_{ab} = 26 * LNC \tag{4}$$

$$C_{ar} = 0.216 * C_{ab} \tag{5}$$

$$LNC = CNC/LAI_{Total} \tag{6}$$

where CNC ($g\ m^{-2}$) represents canopy nitrogen content per unit planting area.

Leaf equivalent water thickness ($C_w$, cm) is an equivalence of leaf water content per leaf area ($g\ cm^{-2}$), referring to the water mass per unit leaf area. It is calculated with Eq (7).

$$C_w = (TLFW - TLDW)/TLA \tag{7}$$

where TLFW (g) is the total leaf fresh weight; TLDW (g) is the total leaf dry weight; TLA ($cm^2$) is the total leaf area. Leaf water content is co-impacted by growth stage, soil water status and weather conditions. In our field experiment conditions, results appear to show that leaf water content was highly correlated with growth stages or fraction of dead leaves at canopy level regardless of cultivar and irrigation-nitrogen application (Figure S1 and Figure S2). Therefore, here $C_w$ was approximately calculated using Eq (8).

$$C_w = \begin{cases} -0.000196\, Z_s + 0.0298 & (f_{dead} = 0) \\ 0.0223 \exp(-1.90\, f_{dead}) & (f_{dead} > 0) \end{cases} \tag{8}$$

where $Z_s$ is the decimal zadok score for the growth stage (Zadoks et al., 1974). The fraction of dead leaves ($f_{dead}$) can be expressed as the ratio of leaf area index of senesced or dead leaves ($LAI_{Dead}$) divided by leaf area index of total leaves ($LAI_{Total}$).

$$f_{dead} = LAI_{Dead}/LAI_{Total} \quad (9)$$

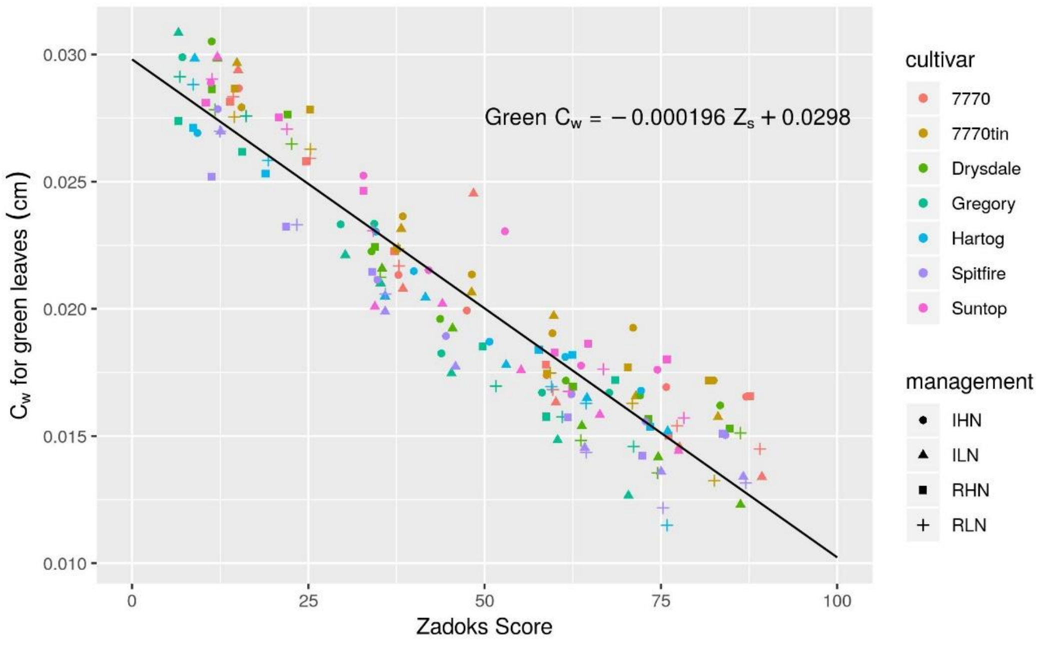


*Figure S1 Leaf water content for green leaves*

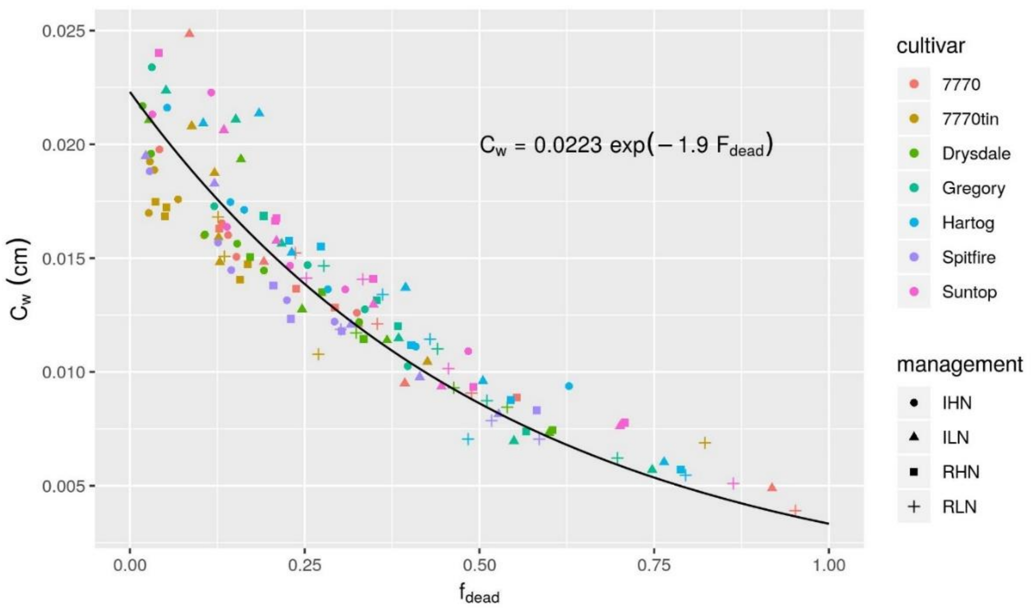


*Figure S2 Leaf water content for total leaves*

The hot spot size parameter (hspot=ι/h) is equal to the ratio of the correlation length of leaf projection in the horizontal plane (ι) and the canopy height (h), which can be estimated in two way (see Verhoef, 1998). The growth of wheat leaf area is mainly achieved by increasing number of leaves, which is usually accompanied by an increase in plant height, so the hspot parameter for wheat can be calculated as:

$$\text{hspot} = a/LAI_{Total} \tag{10}$$

where a is an empirical parameter and here is set as 0.5 (a=0.5) ({Dong, 2019 #539}).

Leaf angle, also called leaf inclination angle, is the angle between leaf surface normal and horizontal surface normal. In nature, leaf angles of individual leaf vary from 0 to 90, but average leaf angles often lie between 20 and 70 (Wilson, 1967). For wheat crops, it is common to have only one or two leaves left on the plant at late growing stage approaching maturity. In this research, based on our knowledge, ALA is set at its common value (50˚) for the aim of simplification. In addition, the output LAI of APSIM-NG can be directly passed to PROSAIL as input variable. The leaf brown pigment parameter ($C_{brown}$) and leaf anthocyanins content ($C_{ant}$) is not considered here and set as zero ($C_{brown}$=0, $C_{ant}$=0) through all growth stages.

As such, nine input parameters of PROSAIL related to crop growth can be converted from APSIM output variables and the remaining four input parameters (SZA, VZA, RAA and $r_{soil}$) associated with environmental conditions for data acquisition are set as shown in Table S2.

*Table S2 Setting for other input parameter of APSIM-PROSAIL used in synthetic dataset generation*

| Parameters | Default | LowerBound | UpperBound | Setting in this research |
|---|---|---|---|---|
| SZA | According to actual condition for collecting data | 0 | 90 | SZA=f(Lat, DOY, t), set t=10:00, 12:00, and 14:00 |
| VZA | | 0 | 90 | Set VZA=0, assuming the sensor's lens face down perpendicular to the ground. |
| RAA | | -180 | 180 | RAA=SAA=f(SZA, DOY, t) when VZA=0. |
| psoil | | 0 | 1 | Set psoil=1, assuming soil surface is dry. |

Notes: t is the apparent solar time (AST); Lat is the latitude; DOY is the day of the year; f denotes a function.

The three sun-sensor-geometry parameters (SZA, VZA and RAA expressed in degree) are determined by the physical environment (location, date and time) where images would be captured. The solar zenith angle (SZA) is the angle between the sun's rays and the vertical panel, which is expressed as the equation below.

$$\cos(SZA) = \sin(L)\sin(\delta) + \cos(L)\cos(\delta)\cos(h) \tag{11}$$

$$\delta = 23.45\sin\left(\frac{360}{365}(284 + DOY)\right) \tag{12}$$

$$h = 15(AST - 12) \tag{13}$$

$$AST = LST + ET \pm (SL - LL) - DS \tag{14}$$

$$ET = 9.87\sin(2B) - 7.73\cos(B) - 1.5\sin(B) \tag{15}$$

$$B = \frac{360}{364}(DOY - 81) \tag{16}$$

where L (°) is local latitude (values of L north of the equator are positive and those south ae negative); δ (°) is the solar declination angle; h (°) is the hour angle; DOY is the day of the year; AST is the apparent solar time (based on the apparent angular motion of the sun across the sky); LST is the local standard time (the time on a clock of a locality); ET (min) is the equation of time; SL (°) and LL (°) is standard and local longitude respectively; DS is the daylight saving (either 0 or 60 min). If a location is east of Greenwich, the sign of Eq. (14) is minus (-), and if it is west, the sign is plus (+). If a daylight-saving time is used, this must be subtracted from the local standard time. The term DS depends on whether daylight saving time is in operation (usually from end of March to end of October) or not. This term is usually ignored from this equation and considered only if the estimation is within the DS period. Here in this research, values of L, DOY and AST are determined by location that crops are growing in and date/time crops are phenotyped. Here L and DOY are set according to APSIM output and, AST is set at three levels (10:00, 12:00 and 14:00) based on practical experience.

The viewing zenith angle (VZA) is the angle between the sensor and the horizontal surface normal where the object is located. It is calculated from coordinates of both sensor and object.

$$\cos(VZA) = \frac{\overrightarrow{OZ} * \overrightarrow{OV}}{|\overrightarrow{OZ}| * |\overrightarrow{OV}|} \quad (17)$$

where $\overrightarrow{OZ}$ is a vector and denotes the vertical direction of object; $\overrightarrow{OV}$ is another vector and denotes the direction from object to sensor. Here the viewing zenith angle is set as zero ($VZA = 0$) as the sensor above the object is placed perpendicular to the object's projection on horizontal surface when imaging.

The relative azimuth angle (RAA) is the difference between solar azimuth angle (SAA) and viewing azimuth angle (VAA). The solar azimuth angle (westward is designated as positive) can be calculated with the equation below.

$$\sin(SAA) = \frac{\cos(\delta)\sin(h)}{\cos(90° - \mathrm{SZA})} \quad (18)$$

If cos(h) > tan(δ)/tan(L), this equation is correct; otherwise, it needs to be adjusted and is (|SAA|-90°) for morning hours and (90°-|SAA|) for afternoon hours.

The viewing azimuth angle is the angle between the sensor and the north direction of object on the horizontal surface. It is obtained from sensor's and object's coordinates.

$$\cos(VAA) = \frac{\overrightarrow{ON} * \overrightarrow{OV^*}}{|\overrightarrow{ON}| * |\overrightarrow{OV^*}|} \quad (19)$$

where $\overrightarrow{ON}$ is a vector and denotes the north direction of object on the horizontal surface; $\overrightarrow{OV^*}$ is another vector and denotes the direction from object to sensor on the horizontal surface. In this case, the viewing azimuth angle needs to be adjusted by minus 180° (that is VAA-180) in order to matching range of solar azimuth angle. Here the viewing azimuth angle is zero as well because the viewing zenith angle is set as zero as mentioned above. Thus, the relative azimuth angle equals the solar azimuth angle.

The reflectance of soil ($r_{soil}$) is determined by the geographical environment where crops grow. In order to simplify the way to illustrate the possibility to adjust soil properties in a continuous fashion, PROSAIL implements a hypothesis that soil reflectance spectrum in general could be represented by mixing a wet soil spectrum and a dry spectrum via adjusting parameter $p_{soil}$. This parameter is a ratio used to mix a dry soil and a wet soil, ranging from 0 (representing a wet soil) to 1 (representing a dry soil). The surface of soil is dry at most time except right after irritation or heavy rain (>5mm). In addition, the influence of soil background can be neglected when LAI>3 according to our sensitivity analysis results and results from (Atzberger et al., 2003) in which situation canopy generally reaches a high ground cover (Ramirez-Garcia et al., 2012). Thus, here in this study it is set as 1 ($p_{soil}$=1).

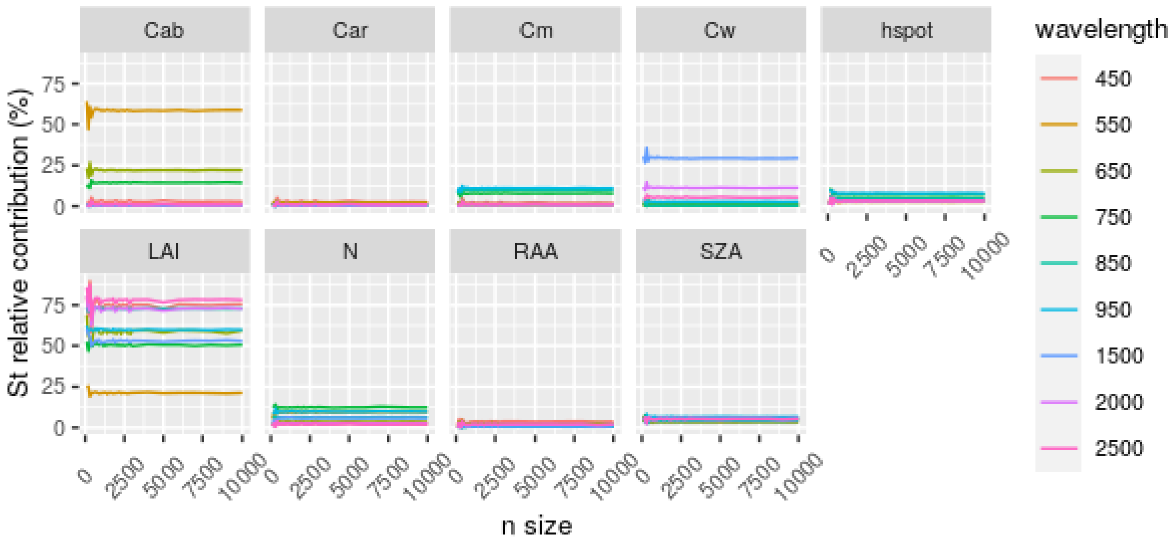


*Figure S3 Relative contribution of total sensitivity index changing with sample size (n) at specific wavelength*

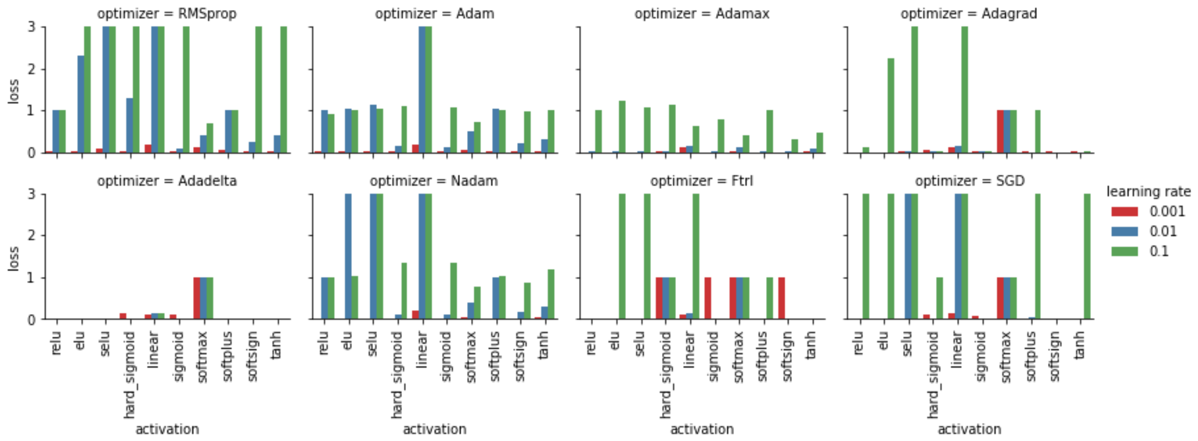


*Figure S4 Loss score of different combinations of learning rate, activation and optimizer. Loss represents the total scaled mean squared error (MSE) of all output variables.*

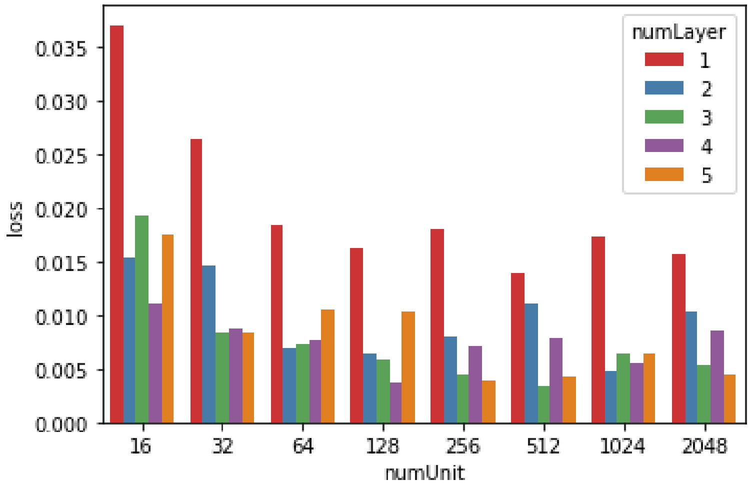


*Figure S5 Loss of different combinations of hidden layer number (numLayer) and unit number (numUnit). Loss represents the total scaled mean squared error (MSE) of all output variables.*

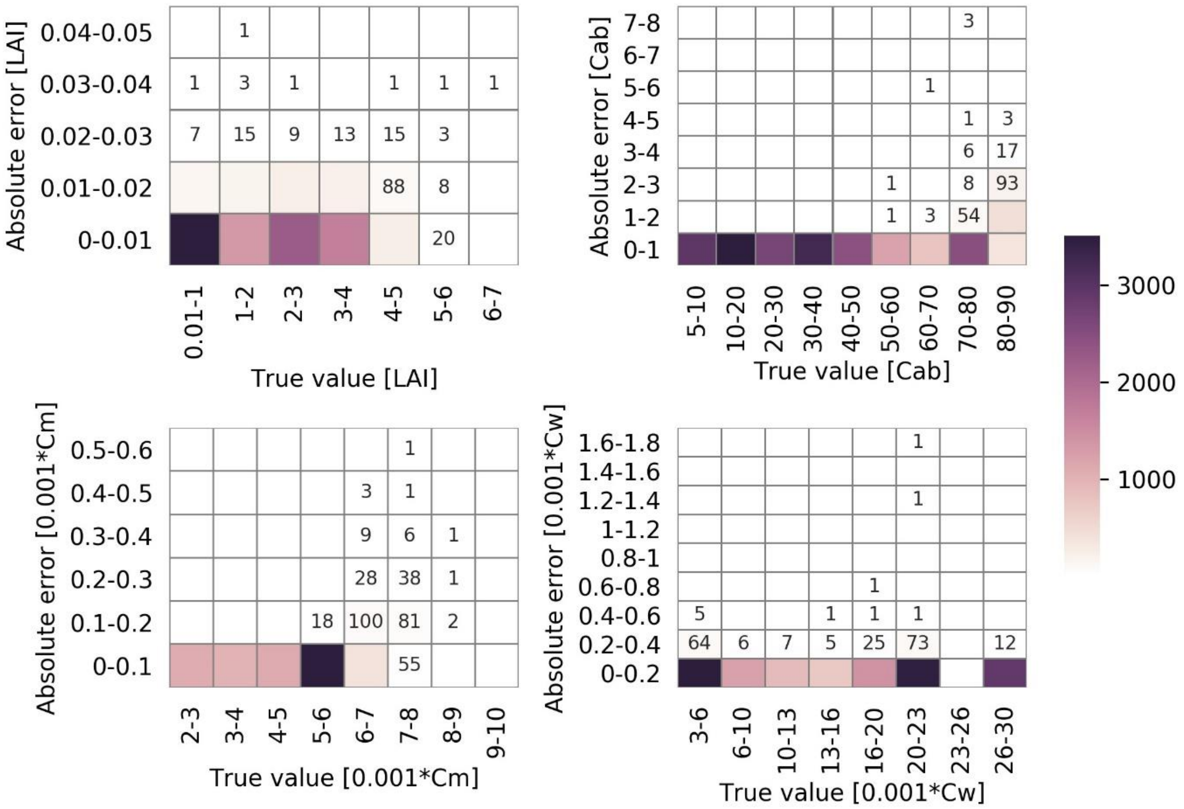


*Figure S6 Heatmap of model performance at different levels (SE8). Numbers in cells of the figure show the number of samples whose prediction absolute errors of true values in a range were within a specific range. Numbers less than 100 are shown to differ small values from zero. The unit of absolute error (true value) of LAI, Cab, Cm and Cw is $m^2$ $m^{-2}$, μg $cm^{-2}$, g $cm^{-2}$ and g $cm^{-2}$, respectively.*

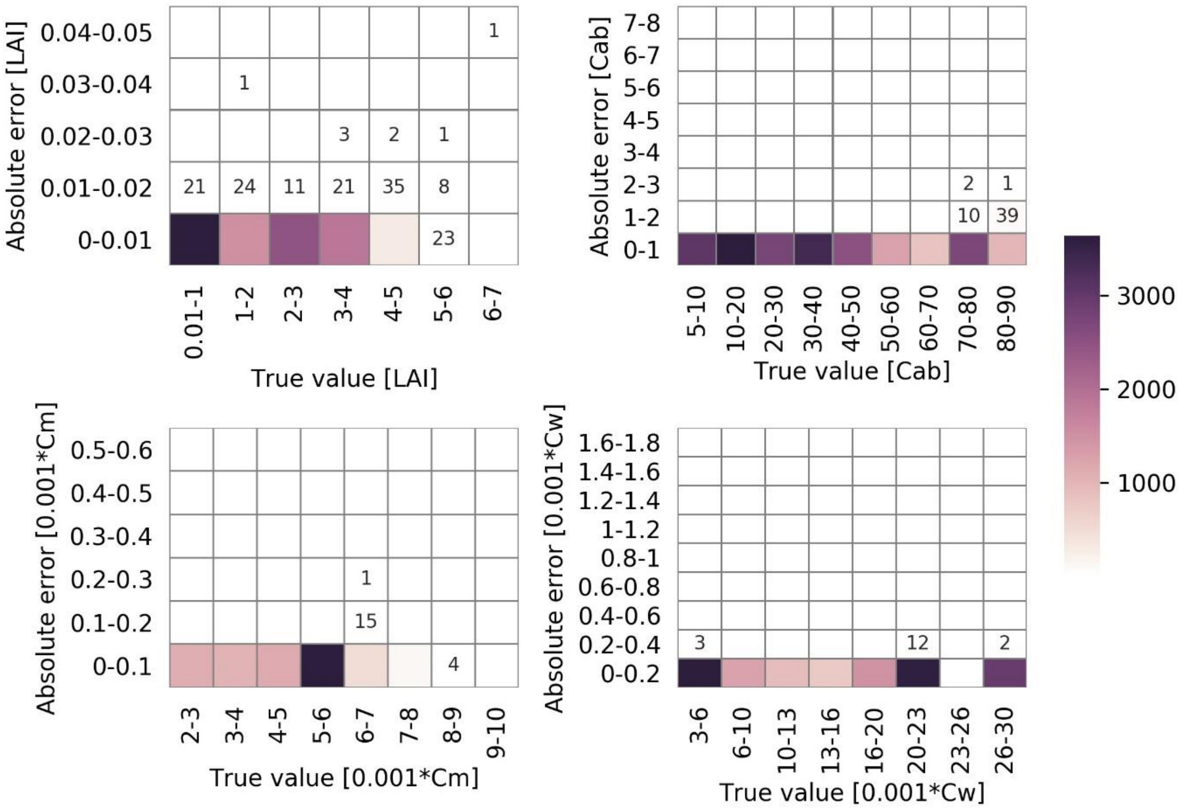


*Figure S7 Heatmap of model performance at different levels (SE9). Numbers in cells of the figure show the number of samples whose prediction absolute errors of true values in a range were within a specific range. Numbers less than 100 are shown to differ small values from zero. The unit of absolute error (true value) of LAI, Cab, Cm and Cw is $m^2$ $m^{-2}$, μg $cm^{-2}$, g $cm^{-2}$ and g $cm^{-2}$, respectively.*
